# Natural diversity of telomere length distributions across 100 *Saccharomyces cerevisiae* strains

**DOI:** 10.1101/2025.05.13.653712

**Authors:** Clotilde Garrido, Cintia Gómez-Muñoz, Etienne Kornobis, Nicolas Agier, Oana Ilioaia, Gilles Fischer, Zhou Xu

## Abstract

Telomeres gradually shorten at each cell division and telomerase counteracts this shortening by elongating telomere sequences. This dynamic balance between elongation and shortening results in a steady-state telomere length (TL) distribution. We developed a method for detecting telomeric sequences in *Saccharomyces cerevisiae* genomes from raw Oxford Nanopore Technologies (ONT) sequencing reads, providing a comprehensive view of TL distributions both genome-wide and at individual chromosome extremities. We analyzed the TL distribution in 100 *S. cerevisiae* strains, representing the genetic and ecological diversity of the species. Our analysis revealed a large diversity in TL distributions within the species, largely driven by inter-extremity differences, ploidy level, and subtelomere structure. Polyploid strains displayed significantly longer telomeres than diploid and haploid strains, and experiments with artificially generated polyploids in two independent genetic backgrounds confirmed that higher ploidy levels lead to telomere elongation. Furthermore, we found that the subtelomeric Y’ element exerts two distinct and opposing effects: (i) the presence of Y’ elements at a chromosome extremity is associated with shorter telomeres in *cis*, but (ii) the overall Y’ element content in a strain correlates with longer telomeres. Interestingly, the length of the shortest telomeres remained relatively constant across strains, suggesting a selective constraint at the species level. This study reveals the diversity of TL in *S. cerevisiae* and highlights key factors shaping TL distributions both genome-wide and at individual chromosome extremities.

## Introduction

Telomeres are repeated sequences found at chromosome extremities, which are essential for genome integrity and control of cell proliferation (de Lange, 2018; Jain and Cooper, 2010; Wellinger and Zakian, 2012). These functions are tightly associated with the regulation of telomere length (TL). Indeed, telomeres are protected by proteins that prevent natural chromosome ends from being recognized as double-strand breaks, which would elicit a DNA damage response and inappropriate repair. In the budding yeast *Saccharomyces cerevisiae*, these include Rap1 and its cofactors Rif1/Rif2, which are bound to double-stranded DNA (dsDNA) in numbers roughly proportional to TL (Levy and Blackburn, 2004; Marcand et al., 1997). In addition, owing to the incomplete replication of chromosome extremities, telomeres shorten at each cell division until they reach a critical length that triggers the DNA damage checkpoint and replicative senescence, a programmed cell cycle arrest that prevents further proliferation (Enomoto et al., 2002; Ijpma and Greider, 2003; Lundblad and Szostak, 1989). TL must therefore maintain a minimal length to allow for cell division. In most species, including budding yeast, this is accomplished by expressing the telomerase enzyme, which preferentially adds telomeric repeats to shorter telomeres (Greider and Blackburn, 1985; Lundblad and Szostak, 1989; Teixeira et al., 2004). In *S. cerevisiae*, telomerase is constitutively expressed, which leads to complex dynamics of progressive telomere shortening interrupted by stochastic elongation by the limiting amount of telomerase that acts on a few telomeres per cell cycle (Gallardo et al., 2011; Mozdy and Cech, 2006; Xu et al., 2013). This length regulation ensures an overall equilibrium, but individual TLs are highly variable across different extremities in a given cell and also across different cells for a given extremity (Shampay and Blackburn, 1988). TL is thus best described by a steady-state TL distribution, and each species appears to have it set at a characteristic length range (Blackburn and Chiou, 1981; Cristofari and Lingner, 2006; Eberhard et al., 2019; Fajkus et al., 1995; Gomes et al., 2011; Monaghan et al., 2018; Moyzis et al., 1988; Shakirov and Shippen, 2004; Walmsley and Petes, 1985; Wicky et al., 1996).

Despite this steady-state TL distribution, the average TL varies broadly across individuals and subpopulations within a species (Allsopp et al., 1992; D’Angiolo et al., 2023; Eberhard et al., 2019; Fulcher et al., 2015; Gatbonton et al., 2006; Karimian et al., 2024; Liti et al., 2009; O’Donnell et al., 2023; Shakirov and Shippen, 2004; Walmsley and Petes, 1985). In *S. cerevisiae*, early studies based on terminal restriction fragment Southern blots revealed moderate variations in the average TL in a limited number of laboratory and wild strains (Gatbonton et al., 2006; Liti et al., 2009; Walmsley and Petes, 1985). More recently, larger variations in TL across natural isolates have been reported (D’Angiolo et al., 2023; O’Donnell et al., 2023). Using short-read whole-genome-sequencing data, D’Angiolo and colleagues found that wild strains display shorter telomeres than strains from domesticated environments, a difference partially driven by mitochondrial metabolism (D’Angiolo et al., 2023). We previously generated telomere-to-telomere genome assemblies for 142 natural yeast isolates (the “*Saccharomyces cerevisiae* Reference Assembly Panel” or ScRAP), which further allowed us to show that TL varies among chromosome-extremities and across strains and that subtelomeric composition contributed to these variations (O’Donnell et al., 2023). Recently, Sholes and colleagues used ONT sequencing to measure the full TL distribution and reported that in the W303 laboratory strain, each chromosome end had a specific and stable TL distribution (Sholes et al., 2022). These observations, since then extended to human telomeres (Karimian et al., 2024; Sanchez et al., 2024; Schmidt et al., 2024), suggest the existence of chromosome-end-specific factors contributing to TL regulation.

Here, we used ONT sequencing to characterize the full TL distribution at each individual chromosome extremity and globally for 100 natural yeast strains. We aim at providing a representative view of TL diversity in *S. cerevisiae* and propose to uncover several features that may underlie TL variations both within and across genomes.

## Results

### Single-molecule sequencing provides a complete view of telomere length distribution

We developed a pipeline to identify telomeric sequences in individual ONT reads (Fig. 1A). After base calling and adapter removal, the reads are mapped onto the genome assembly of the target strain. The reads that align with chromosome extremities are scanned for telomeric sequences using Telofinder, a script designed to specifically detect the degenerated yeast telomere motif TG_1-3_/C_1-3_A (O’Donnell et al., 2023). Finally, after filtering steps ensuring that the telomere sequences are not truncated or internal (Supp. Fig. S1A), the lengths of all validated telomere sequences are aggregated into a high-resolution TL distribution (Fig. 1B-C). Importantly, because typical ONT reads are tens to hundreds of times longer than TL, the read length did not affect the computed TL distribution (Supp. Fig. S1B), ensuring the robustness of the method with respect to variations during DNA extraction and sequencing.

**Figure 1.**
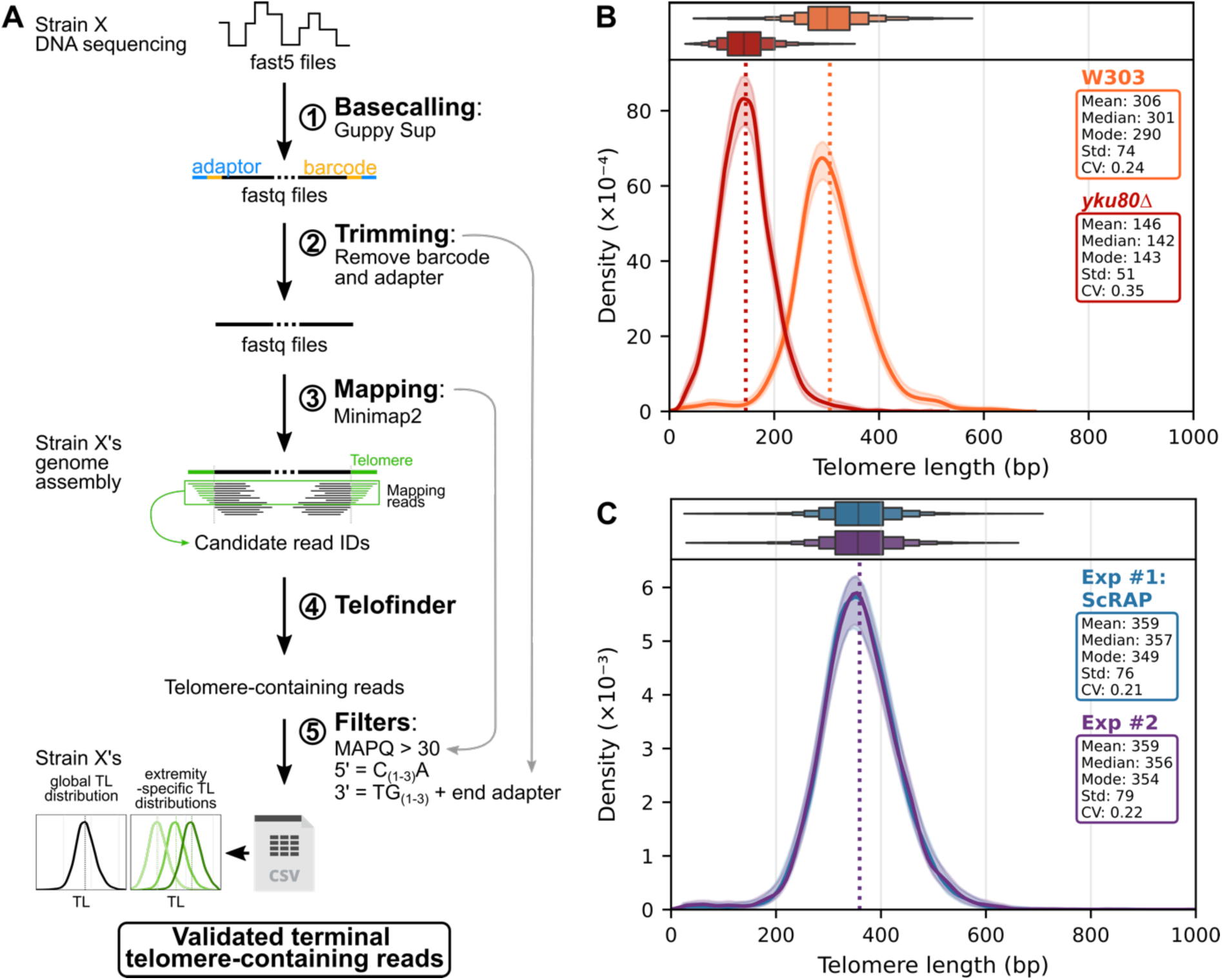
High-resolution TL distribution from ONT sequencing. (A) Bioinformatic pipeline to retrieve TLs from single-molecule ONT sequencing. The raw electric signal from nanopores was basecalled using the super accuracy mode of Guppy (step 1). Barcodes and adapters were then removed by Porechop (step 2) and candidate sequences mapping to the telomere regions of the corresponding genome assembly were selected (step 3). The initial reads corresponding to these candidates were scanned for telomere sequences using Telofinder (step 4) and validated as telomeric reads if compliant with specific quality filters (step 5), including a mapping quality (MAPQ) threshold and a test for the presence of an adapter at the 3’ end of the molecule for TG_(1-3)_ sequences (gray arrows). Global and extremity-specific TL distributions are then computed. (B) TL distributions of the wild-type W303 strain and its *yku80Δ* derivative, represented by a boxplot (upper graph; successive box edges indicate 50%, 75%, 87.5%, 93.75%, etc., of the data) or by its density (lower graph), obtained from ONT sequencing using the bioinformatic analysis pipeline described in (A). The shaded area indicates the 95% confidence interval of the sampling noise, inferred from bootstrapping. The mean value is indicated by the dotted line. The detailed metrics are shown in the insets. (C) TL distribution of strain AHG, using data from (O’Donnell et al., 2023) (“Exp #1”) and from a new sequencing (“Exp #2”). Same representation as in (B).

First, we tested the ability of our methodology to detect biologically relevant TL differences by comparing TL distributions between the wild-type (WT) W303 laboratory strain and its *yku80Δ* derivative, in which the function of the Ku complex is impaired, leading to significantly shorter telomeres (Porter et al., 1996). Our method measured a mean TL that was ∼164 bp shorter in *yku80Δ* compared to WT (Fig. 1B and Supp. Fig. S1C and D), which is fully consistent with previously reported estimates (Bertuch et Lundblad 2003; Chen et al. 2018). The TL distribution in the *yku80Δ* mutant was also narrower (average standard deviation of 60 bp compared with 77 bp in the WT), likely owing to constraints preventing telomeres from becoming too short.

Second, we tested the reproducibility of our methodology by comparing TL distributions between independent sequencing experiments derived from biological replicates. We used a natural strain isolated from pear must (AHG) for which the first set of reads was recovered from the ScRAP study (O’Donnell et al., 2023), and the second set of reads was obtained from a new ONT sequencing performed in the frame of this study. We found an excellent agreement between the results of 2 independent experiments (Fig. 1C). We also tested the robustness of TL distribution measurements across ONT technologies and sequenced biological replicates of the W303 strain and its *yku80Δ* derivative with both R9.4.1 flowcells and the currently available R10.4.1 flowcells. The two flowcell technologies yielded nearly identical distributions (Supp. Fig. S1C-E). Nonetheless, for the sequencing based on the R9.4.1 flowcells, the comparison between TL distributions derived from the C-rich strand and the G-rich strand showed a minor difference in means (5-20 bp longer for the G-rich distribution), which was completely eliminated with the R10.4.1 flowcells (Supp. Fig. S1E), indicating a slight technical, rather than biological, bias associated with R9.4.1 flowcells.

Overall, our method derives accurate, robust and biologically informative TL distributions directly from ONT sequencing reads. It is highly versatile, as it can be applied to any *S. cerevisiae* sequencing dataset without any particular telomere enrichment step, enabling the investigation of telomere length distribution diversity across numerous strains.

### TL distributions are highly diverse across *S. cerevisiae* strains

We measured TL distributions in a panel of 100 strains, representing the ecological and geographical diversity of *S. cerevisiae* (O’Donnell et al., 2023) (Fig. 2A and B). This representative dataset provided a general overview of TL distribution at the species level by aggregating telomeric reads from all 100 strains (Supp. Fig. S2A). The mean ± standard deviation TL was measured at 366 ± 137 bp, which is slightly longer than commonly reported estimates in laboratory strains (Wellinger and Zakian, 2012), with the 1^st^ and 99^th^ TL percentiles at 112 and 827 bp, respectively.

**Figure 2.**
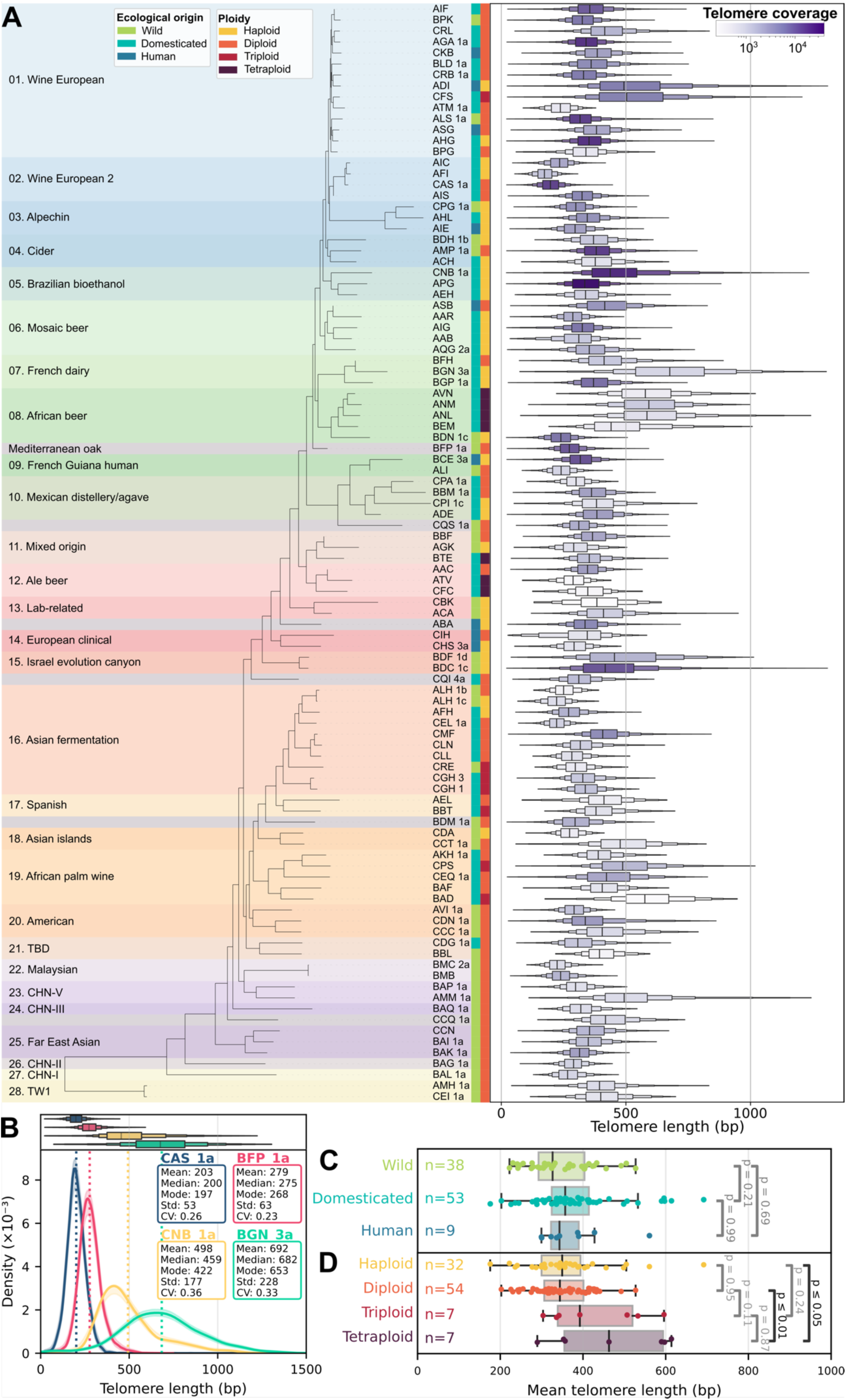
Natural diversity of the TL distribution in *S. cerevisiae*. (A) Phylogenetic tree and clade classification from (O’Donnell et al., 2023). For each strain, the TL distribution is represented as a boxplot. (B) TL distributions for 4 strains, selected to illustrate the diversity in means and standard deviations. (C) Boxplots of the TL means of wild, domesticated and human-associated strains. *p*-values from Tukey’s HSD tests are shown (initial ANOVA test *p*-value = 0.22). (D) Boxplots of TL means of haploid, diploid, triploid and tetraploid strains. *p*-values from Tukey’s HSD tests are shown (initial ANOVA test *p*-value = 0.003). Non-significant differences are shown in gray and significant ones in black.

Among the strains, the mean TL varied significantly, ranging from 176 bp in strain AFI, associated with European wine, to 682 bp in strain BGN_3a, sampled from French dairy, with ∼50% of strains having TL means within the 300-400 bp range. Examples of 4 strains with diverse TL distributions in terms of means and standard deviations are shown in Fig. 2B. Within the same clade, while some strains presented similar TL distributions (e.g., ADI and CFS, from clade “Wine European”), others exhibited vastly different ones (e.g., BAP_1a and AMM_1a, from clade “CHN-V”) (Fig. 2A and Supp. Fig. S2B-D). Notably, differences between mean TLs did not correlate with phylogenetic distances, whether within or between clades (Supp. Fig. S2B). A correlation could only be detected for the most closely related strains (R² = 0.4 for the top 4% closest pairs of strains, Supp. Fig. S2C). Overall, this level of TL variation across strains far exceeded that observed between biological replicates (Fig. 1C, Supp. Fig. S1C and D), indicating that TL is controlled in a strain-specific manner.

Additionally, comparing TL distributions across strains revealed striking variations in other metrics, including several that were previously inaccessible. Notably, the standard deviation of the distributions varied from 44 to 228 bp, correlating with the mean (Supp. Fig. S2E). The coefficient of variation (CV, computed as the standard deviation divided by the mean) was around 0.26, but varied from 0.19 to 0.39, indicating that the standard deviation did not simply scale with the mean TL (Supp. Fig. S2F). Skewness displayed values from −0.51 to 4.79, with most strains (n = 85 out of 100) exhibiting positively skewed distributions, an asymmetry consistent with the counterselection of critically short telomeres, as supported by mathematical modelling (Xu et al., 2013). The robust kurtosis values ranged from 3.41 to 7.22, with a mean of 5.22, indicating a tendency toward leptokurtic distributions, i.e. shaped with sharper peaks and heavier tails than a normal distribution, suggesting a rather loose regulation of very long telomeres.

The factors that control TL distributions and their variations across strains are not known. We tested whether the ecological origins of the strains could influence TL but did not find any significant difference in the mean TL (Fig. 2C). Although not significant in our results possibly because of the limited sample size, there was a trend toward longer telomeres in domesticated strains than in wild strains (Fig. 2C), which is consistent with recent findings (D’Angiolo et al., 2023).

### Polyploidy leads to longer telomeres

We found that natural polyploids, particularly tetraploids, exhibited significantly longer mean TLs than haploids and diploids did (Fig. 2D), a trend that had been observed, although not statistically significant, in our previous study (O’Donnell et al., 2023). To experimentally test the effect of polyploidy on TL, we generated homozygous tetraploid strains by recursively crossing mating-proficient versions of 2 independent homozygous natural diploid strains (YPS128 from North America and DBVPG6044 from West Africa) with their haploid derivative (Supp. Fig. S3A) (Cubillos et al., 2009; Louvel et al., 2014). Strikingly, in both backgrounds, the tetraploid strain displayed a global TL mean substantially longer than the TL mean of the initial diploid strain (391 ± 83 bp compared to 324 ± 66 bp in YPS128 background; 453 ± 98 bp compared to 415 ± 96 bp in DBVPG6044 background) (Fig. 3A and B), and longer than the TL mean of the haploid derivatives (Supp. S3B and C).

**Figure 3.**
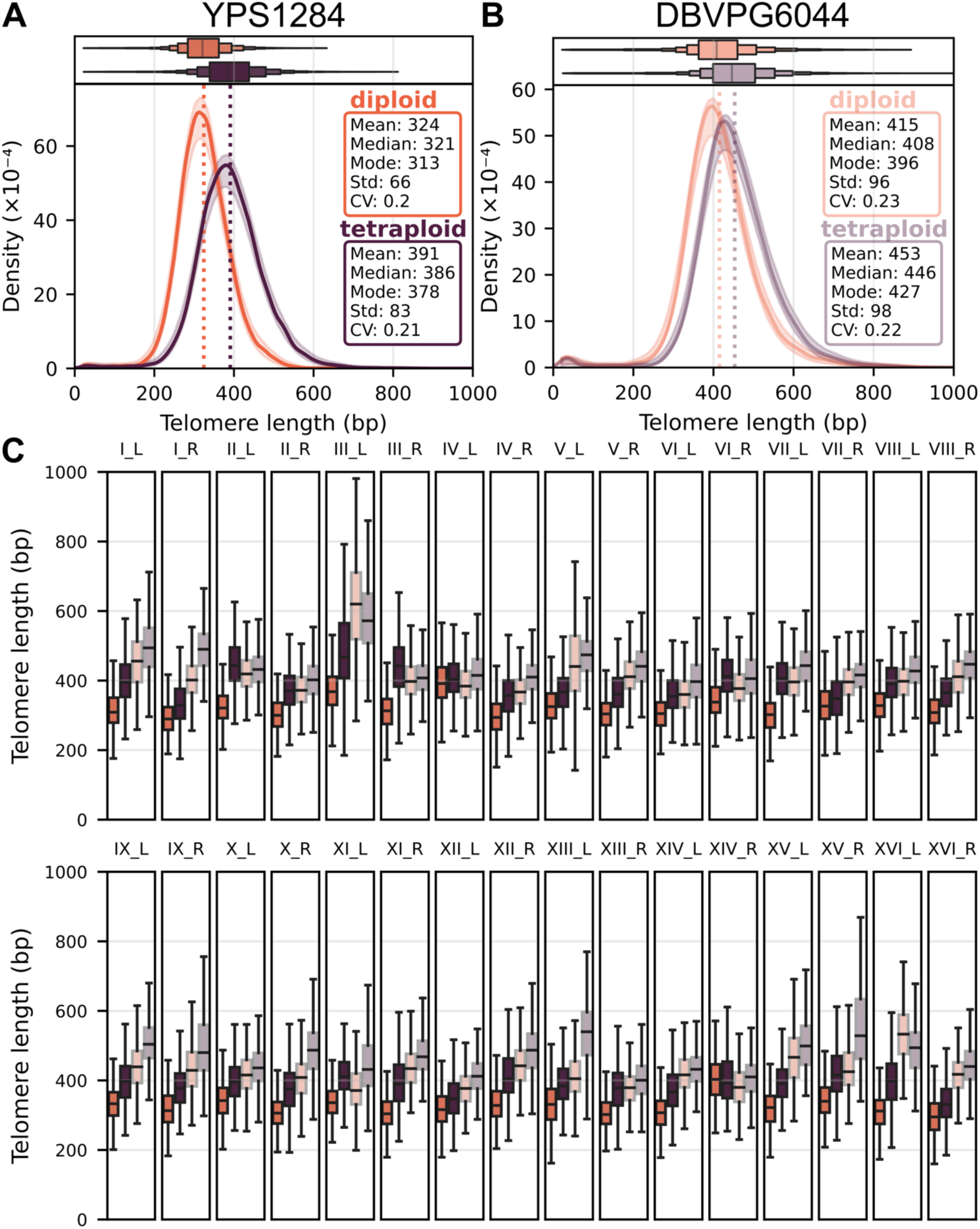
Tetraploids exhibit longer telomeres than isogenic homozygous diploids. (A) Global TL distributions of the natural North American diploid strain YPS128 and the experimentally generated isogenic tetraploid. (B) Global TL distributions of the natural West African diploid strain DBVPG6044 and the experimentally generated isogenic tetraploid. (C) Boxplots of extremity-specific TL distributions for all extremities, shown for the 4 strains from (A) and (B) (same color code).

In addition, by assigning each telomere-containing read to a specific chromosome end using the subtelomere sequence as an anchor, our pipeline was also able to derive the complete distributions of extremity-specific TLs (Fig. 3C). The TL difference between the tetraploid and diploid strains was observed at 31 and 30 out of 32 individual chromosome extremities, for YPS128 and DBVPG6044 backgrounds, respectively, indicating that TL elongation in the tetraploid strains reflects a genome-wide regulatory effect rather than changes at specific chromosome extremities (Fig. 3C). We thus conclude that ploidy increase beyond diploidy leads to longer telomeres.

### TL distributions are extremity-specific and inter-extremity differences vary across strains

Chromosome-end-specific differences in steady-state TL have been reported in laboratory yeast strains, using ONT sequencing (Sholes et al., 2022). We previously extended this observation to diverse natural yeast strains using estimates based on genomes assemblies (O’Donnell et al., 2023). These observations were enabled by the long reads allowing accurate anchoring of telomere sequences to their corresponding chromosome extremity.

Here, beyond deriving strain-specific TL distributions, we obtained the distributions of extremity-specific TLs (Fig. 4A). The means of these extremity-specific TL distributions correlated well with the TL values obtained from the genome assemblies (r^2^ = 0.9), with most values differing by less than 10% (Supp. Fig. S4A). However, for 12% (resp. 1%) of the extremities, the assembly-based TL values underestimated (resp. overestimated) the mean TL by 20% or more. Importantly, the method presented in this work not only corrects for the assembly-level inaccuracies at telomeres but also provides whole extremity-specific TL distributions, which are not accessible from genome assemblies and contain much richer information.

**Figure 4.**
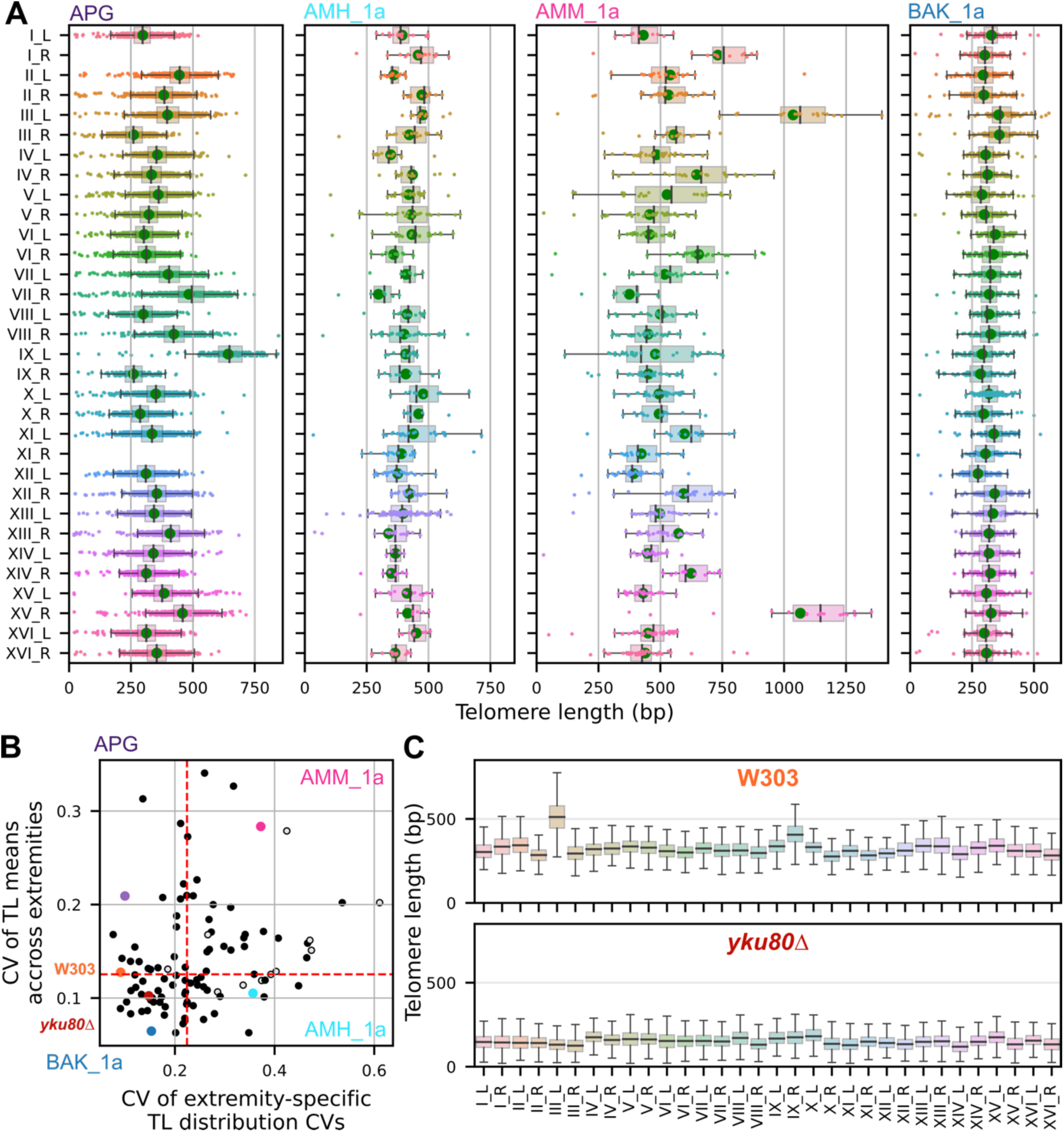
Diversity of extremity-specific telomere length in natural *S. cerevisiae* strains. (A) TL distributions at every chromosome extremity for 4 selected strains, represented as boxplots with individual read TLs plotted as dots. (B) 2D plot displaying the CVs of extremity-specific TL means and the CVs of extremity-specific TL distributions CVs. Each strain is represented as a dot and this 2D space is divided vertically and horizontally into 2 equally sized areas. Open dots indicate strains for which the sample size of the reads was insufficient for confident inference. The 4 strains shown in (A) as well as W303 and its *yku80Δ* derivative are labelled in color. (C) TL distributions at every chromosome extremity for W303 and its derived *yku80Δ* mutant.

While we confirmed that TL means were extremity-specific for most strains, we found that the extent of the inter-extremity differences within a strain varied widely (Fig. 4A). For example, strain APG associated with Brazilian bioethanol exhibited differences in mean TL between extremities whereas the standard deviation across extremities remained rather constant. Conversely, strain AMH_1a (clade “TW1”) presented variable standard deviations but quite homogeneous TL means. In contrast, in AMM_1a (clade “CHN-V”), both the TL means and standard deviations (and even CVs) were highly variable from extremity to extremity. Finally, other strains displayed similar extremity-specific TL distributions in terms of both the mean and standard deviation at all extremities (e.g. BAK_1a, from clade “Far East Asian”). To systematically survey the inter-extremity differences both in means and CVs (as illustrated by the 4 examples in Fig. 4A), we plotted the inter-extremity CV of the means and the inter-extremity CV of the CVs on a 2D graph for all strains (Fig. 4B). Here, the CV was chosen to measure variation preferentially over the standard deviation, because it allows direct comparison by normalizing the standard deviation by the mean. We took into account the possibility that, for some strains, the coverage by telomere-containing reads might be too low to confidently derive the inter-extremity CV of the means or CV of the CVS, and estimated the minimum coverage required for each strain (Supp. Fig. S4B-K). The coverage of 11 strains did not meet the criteria and their inter-extremity CV of the mean or CV of the CVS were likely overestimated (Fig. 4B, open dots, and example in Supp. Fig. S4B and C).

Our panel of strains was spread over the 2D space of this plot, revealing distinct TL distribution patterns across extremities. We also placed W303 and its derived *yku80Δ* mutant on this plot, which revealed that they presented low inter-extremity variations (Fig. 4B). Upon closer inspection of their extremity-specific TL distributions, we found that the inter-extremity differences displayed in the wild-type W303 were suppressed in *yku80Δ*, particularly the longer TL at ChrIII_L (Fig. 4C). Overall, strains can exhibit high variability in extremity-specific mean TL or CV, or both, suggesting different drivers of TL diversity within the same species.

### Large global TL variations primarily result from inter-extremity differences

Given the substantial inter-extremity differences in mean TL observed at the strain level, we systematically investigated their contribution to global TL variations compared with the contribution of within-extremity variations. Put differently, do global strain-specific TL variations primarily reflect TL variations at each extremity or are they driven by differences in the mean TL between extremities?

To quantify the contribution of inter-extremity differences for each strain, we computed the CV of the extremity-specific TL means, reflecting the inter-extremity variation, and plotted it against the CV of the global TL distribution (Fig. 5A, red, and 5B). We found a strong correlation between these two values across strains (r^2^ = 0.82, p-value = 4×10^-25^). Strikingly, the slope of the correlation was equal to 1.1, indicating that any increase in global TL variation stemmed mostly from accrued inter-extremity differences. Conversely, we plotted the mean of the CVs of extremity-specific TL distributions in a strain, reflecting the average within-extremity variation, against the CV of the global TL distribution (Fig. 5A, blue). We observed that the mean extremity-specific CV scaled with the global CV, with a much lower slope of 0.3 and correlation (r^2^ = 0.53, p-value = 1×10^-8^). Even strains with high global CVs (e.g. >0.3) still showed moderately increased extremity-specific CVs of 0.2-0.25, differing only slightly from strains with lower global CVs.

**Figure 5.**
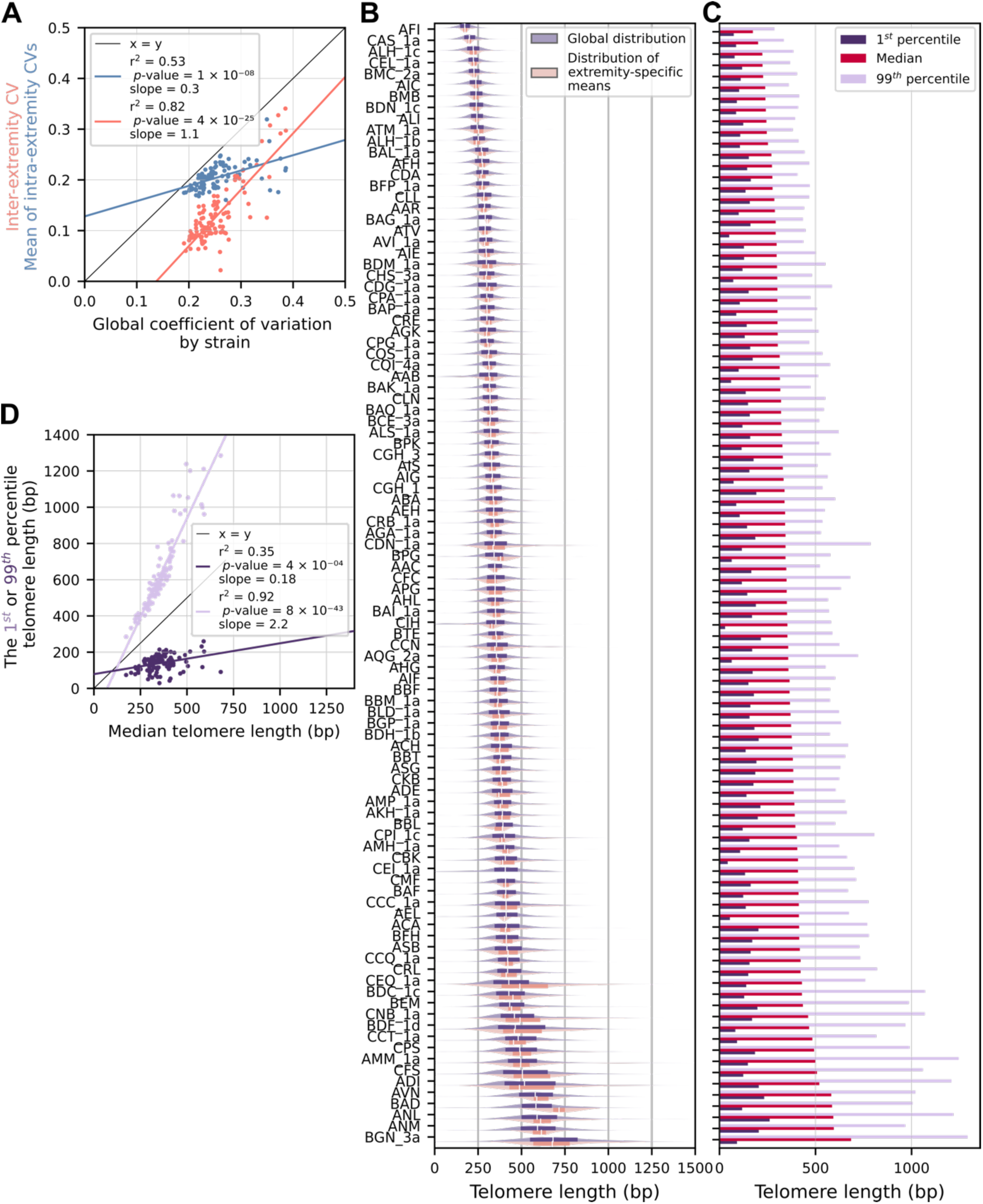
Contributions to global TL variations. (A) The mean of extremity-specific TL distribution CVs, reflecting the average within-extremity variation, is plotted against the global TL distribution CV as a blue dot for each strain. The CV of the distribution of extremity-specific TL means, reflecting inter-extremity variation, is plotted against the global TL distribution CV as a red dot for each strain. (B) Global TL distribution (blue, upper part of each graph) and distribution of extremity-specific TL means (salmon, lower part of each graph) are displayed for all strains, ranked by increasing median TL. A boxplot representing 50% of the values is superimposed on the density plot. (C) Median, 1^st^ and 99^th^ percentiles of the global TL distribution are displayed for each strain. Strains are ordered by increasing median TL. (D) The 1^st^ (purple) and 99^th^ (mauve) percentiles of the global TL distribution for each strain are plotted against the median TL and the linear regressions are shown.

Thus, global TL distribution variation at the strain level comprises two components: the variations found at each extremity-specific TL distribution and the differences in TL between extremities. The latter accounts for most of the observed TL variations between strains.

### The length of the shortest telomeres is constant at the species level

Telomerase acts preferentially on shorter telomeres and prevents them from reaching a critical length (Teixeira et al., 2004). Telomere elongation, combined with progressive telomere shortening, thus constrains the average length of the shortest telomeres in a population of cells. Given the extent of natural TL variations that we revealed, we wondered whether different strains presented different lengths for the shortest telomeres.

We thus calculated the lowest percentile of the TL distribution of each strain and compared it to the median TL (Fig. 5C). When plotted for all strains ranked according to increasing median TL, we found that the lowest percentile did not increase substantially and fluctuated around 138±45 bp, whereas the 99^th^ percentile (longest telomeres) did. To quantitatively evaluate this observation, we calculated the Pearson’s correlation values of the 1^st^ and 99^th^ percentiles with the median and obtained correlations of r^2^ = 0.35 and 0.92, respectively (Fig. 5D). The value of the 1^st^ percentile was thus only marginally correlated with the median TL length across strains.

To assess the extent to which the length of short telomeres was poorly correlated with the median, we plotted the Pearson’s correlation between the n^th^ percentile and the median TL and found that the 2^nd^ (resp. 3^rd^) percentile had a r^2^ value of 0.66 (resp. 0.81), which was significantly lower than the plateau at 1 (Supp. Fig. S5A). Interestingly, for a haploid genome with 32 telomeres, the length of the shortest telomere should be found among the 1/32 = 3.1% lowest TL values in a distribution. The relatively weak correlation between the 1^st^-3^rd^ percentiles and the median TL suggested that the length of the shortest telomere might be kept constant at the species level, regardless of the median TL.

Since extremity-specific differences in TL are observed in many strains, we wondered whether some particular chromosome extremities might bear shorter TLs more frequently than others do. We thus evaluated the representation of each extremity in the individual telomeres shorter than the 1^st^ (resp. 5^th^) percentile across strains and found no significant enrichment of any of them (Supp. Fig. S5B, resp. S5C), indicating that the telomere of any chromosome extremity could be among the shortest. In contrast, we found an overrepresentation of ChrIII_L, ChrXII_R, ChrXIII_R and ChrXV_R among telomeres longer than the 99^th^ percentile (only ChrIII_L among telomeres longer than the 95^th^ percentile) of the distributions across strains (Supp. Fig. S5D and S5E). To perform further analysis, we then asked whether the shortest extremity-specific telomere on average in a strain would be enriched among the individual telomeres shorter than the 1^st^ (resp. 5^th^) percentile (Supp. Fig. S5F, resp. S5G). For each strain, we ranked each chromosome extremity from the shortest associated mean TL to the longest associated mean TL and calculated their enrichment within the 1^st^ (resp. 5^th^) percentile across strains. Even here, we barely found an enrichment for the strain-specific extremity associated with the shortest and second shortest mean TLs within the 1^st^ percentile (resp. the 5 shortest TLs were enriched within the 5^th^ percentile), with Log2 enrichment values below 1 (resp. 2) (Supp. Fig. S5F, resp. S5G). Conversely, extremities associated with greater mean TL in each strain were highly overrepresented within the 99^th^ (resp. 95^th^) percentile or longer with Log2 enrichment values exceeding 4 (resp. 2) (Supp. Fig. S5H, resp. S5I). Overall, we demonstrate that throughout the diversity of natural yeast strains, any chromosome extremity can bear the shortest telomere(s) with nearly equal frequency, an important finding to consider when studying of the functional consequences of the shortest telomere(s).

### Subtelomeric Y’ and Ty5 elements are associated with telomere length variation in *cis*

The observation that TL distributions are often extremity-specific suggested the presence of *cis* regulatory elements. We thus compared extremity-specific TL distributions across strains, with or without elements commonly found at subtelomeres such as X and Y’ elements, and Ty5 retrotransposons. To account for different global TL means in different strains, we normalized each extremity-specific TL by the global TL mean of the corresponding strain, thus allowing the aggregation of normalized values across strains. The presence or absence of the X element in the adjacent subtelomere did not significantly affect the normalized extremity-specific mean TL (Fig. 6A). In contrast, the presence of Ty5 in the *cis* subtelomere was associated with significantly greater normalized extremity-specific mean TL (Fig. 6B), although the imbalance between the number of subtelomeres with and without Ty5 prevented more detailed analyses at the level of each extremity across strains.

**Figure 6.**
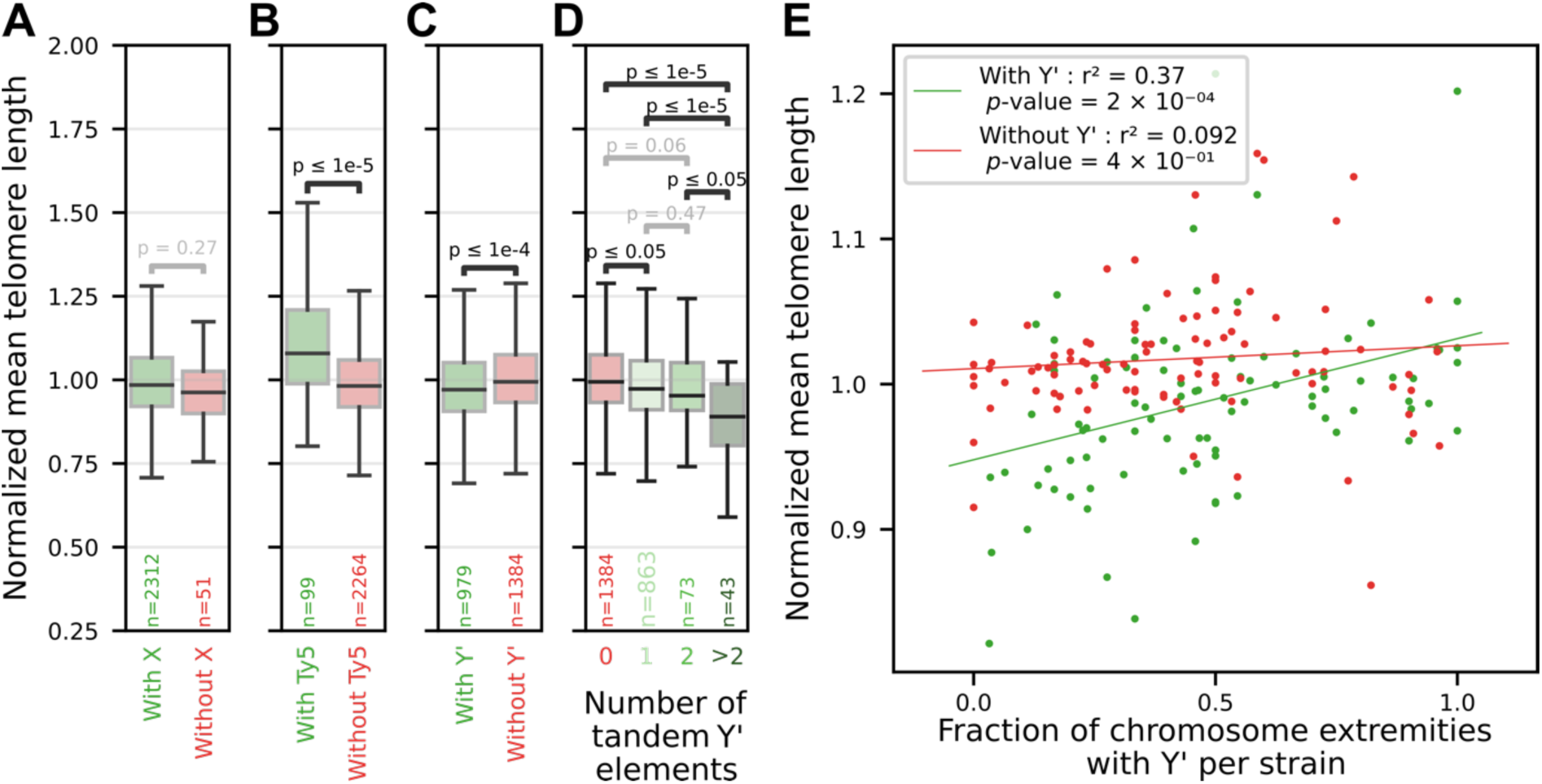
Association between the presence of subtelomeric elements and telomere length. (A) Distributions of the normalized extremity-specific TL means across the extremities of all strains, clustered according to the presence or absence of the X element in the corresponding subtelomere. The number of chromosome extremities in each group is indicated by “n”. The *p*-value from a Student’s t-test is indicated. (B) Same as (A) but with respect to the presence or absence of the Ty5 retrotransposon in the corresponding subtelomere. (C) Same as (A) with respect to the presence or absence of Y’ elements in the corresponding subtelomere. (D) Same as (C) but with respect to the number of tandem Y’ elements in the corresponding subtelomere. (E) For each strain, extremities are clustered according to the presence (green dots) or absence (red dots) of Y’ element. For each strain and within each cluster, the average of normalized extremity-specific TLs is plotted against fraction of extremities with Y’ elements in the same strain.

The presence of Y’ elements in the subtelomere was associated with a significantly lower normalized extremity-specific mean TL (Fig. 6C), as well as increased standard deviation, CV and skewness (Supp. Fig. S6A-C). This effect was significantly stronger for chromosome extremities bearing more than 2 Y’ elements in tandem (Fig. 6D). Looking closely at each chromosome extremity across strains, this negative association between presence of Y’ elements and TL was verified for 21 out of 32 extremities (66%), and reversed only at 4 extremities (Supp. Fig. S6D).

Since interstitial telomere sequences (ITS) are often found centromere-proximal to Y’ elements, they could act as a likely negative regulator of TL in *cis*, but we did not detect a significant correlation between the normalized extremity-specific mean TL as a function of total ITS length in the corresponding subtelomere (Supp. Fig. S6E). Interestingly, without normalization by the global TL mean per strain, the relationship was reversed and the presence of Y’ elements at a chromosome extremity appeared rather associated with longer telomeres *in cis* (Supp. Fig. S6F). This was due to a positive correlation between the global TL mean per strain and the fraction of subtelomeres bearing Y’ element in each strain (Supp. Fig. S6G). When the chromosome extremities of each strain were clustered into Y’-containing and non-Y’-containing, we found that the mean TL of Y’-containing extremities in a strain increased as a function of the fraction of Y’-bearing extremities in the same strain (Fig. 6E). In contrast, the mean TL of non-Y’-containing extremities did not correlate with this fraction. Thus, Y’ elements had two distinct effects on TL: (i) TL positively correlated with the overall Y’ content in a strain, but only for chromosome extremities with Y’ elements, revealing a positive *trans* effect, and (ii) within each strain, Y’-associated telomeres were, however, shorter than telomeres at other extremities, indicative of a negative *cis* effect.

Overall, Y’ elements might provide a local genomic feature as a cause for inter-extremity TL differences, both through their *trans* and *cis* effects. Whether the Y’ element directly regulates TL through a binding factor or alters the local chromatin structure thereby affecting TL homeostasis, remains to be investigated.

## Discussion

*S. cerevisiae* is a major genetic model organism in the telomere field (Teixeira, 2013; Wellinger and Zakian, 2012). In contrast to the large body of work using laboratory strains, relatively little is known about TL regulation and distribution in natural yeast strains. Recent large-scale population genomics studies revealed great genetic diversity encompassing, beyond single nucleotide polymorphisms, structural variations, aneuploidies and polyploidies (O’Donnell et al., 2023; Peter et al., 2018). We consider TL as yet another genomic variant, albeit one that is dynamic and complex, but its diversity within the species has not yet been investigated in depth. Exploring and characterizing TL distributions in natural strains would expand our understanding of telomere regulation and help interpret telomere biology in light of the budding yeast natural history, ecology and evolution. As a first step toward this goal, in this work, we provide a detailed characterization of TL distributions in a panel of natural isolates recently sequenced by long-read ONT sequencing (O’Donnell et al., 2023) and derive biologically relevant insights.

We observed a large diversity of TL distributions across a representative panel of strains. Compared with the commonly studied laboratory strains, some strains present much shorter or longer mean TLs, which might be explained by genetic variants and differences in the molecular mechanisms governing TL regulation. Genome-wide association studies (GWAS) or linkage analyses could leverage the genetic diversity to identify new genes or variants contributing to TL regulation, an approach which has yielded important insight into telomere length regulators (Gatbonton et al., 2006; Liti et al., 2009). Furthermore, TL diversity goes beyond variations of the mean. The standard deviation, CV, skewness and kurtosis all show substantial differences, even within the same clade. Because TL distribution is fundamentally shaped by the dynamics of telomere shortening and elongation by telomerase, the diversity of TL distributions in natural strains suggests that they differ in some or all aspects of these mechanisms.

For example, in our dataset, natural polyploids exhibit longer telomeres than diploids and because domesticated strains are particularly enriched in polyploids (Supp. Table S1), this may partially account for their tendency to have longer telomeres (D’Angiolo et al., 2023). Notably, we demonstrate that experimentally generated tetraploid strains display longer telomeres than the isogenic natural diploid strains, across virtually all chromosome extremities, revealing a causal link between ploidy and telomere length regulation.

For ChrIII-L specifically, Teplitz and colleagues proposed a molecular mechanism involving heterochromatin on the HML locus promoting telomerase recruitment through the Sir4-yKu pathway in the absence of the Y’ element at the ChrIII-L subtelomere to explain the longer telomeres observed without the Y’ element at this extremity (Teplitz et al., 2024). Consistently with their results, we found that inter-extremity TL differences, including longer ChrIII-L telomeres, are eliminated in the *yku80Δ* mutant compared with wild-type W303, implicating the Ku complex in mediating TL regulation in cis. However, another study showed that deleting all X elements, Y’ elements or both in a strain engineered to contain only 3 chromosomes had no significant effect on TL, as measured by Southern blot (Hu et al., 2024), although this technique might not have the resolution to detect subtle differences.

We leveraged the structural diversity found at the subtelomeres of the ScRAP panel of strains to investigate the contribution of subtelomeric elements in *cis* to TL regulation. We detected a small but significant decrease in the normalized extremity-specific TL mean at chromosome extremities containing one or more Y’ elements. Closer investigation shows that Y’ elements exert their effects at two levels: (i) a positive *trans* effect whereby the overall Y’ content is associated with longer TL at Y’-containing extremities, and (ii) a negative *cis* effect whereby the presence of Y’ elements is associated with shorter TL at the corresponding chromosome end. We thus propose that, as a general rule, Y’ elements and possibly Ty5 retrotransposons, but not X elements, influence TL in *cis*, thus contributing to TL variations between extremities. A plausible mechanism to explain the complex relationship with Y’ elements would be the involvement of a Y’-binding factor that negatively regulates TL in *cis*, and the strength of this regulation would weaken if more Y’ elements are present in the genome due to dilution or titration of the limited pool of this factor, thus leading to an apparent positive regulation at a global level. While this putative factor is not known, it is not related to the ITS commonly found at the centromere-proximal side of the Y’ elements. Further investigations are needed to fully elucidate the molecular mechanisms controlling this aspect of TL regulation.

While yeast cells constitutively express telomerase, the signaling pathway triggering replicative senescence if a telomere becomes critically short is functional, as revealed in telomerase mutants (Lundblad and Szostak, 1989). Selection against critically short telomeres could therefore shape the regulation of TLs so that, with active telomerase, the steady-state TL distribution would not allow senescence to occur at a significant frequency. Conversely, cells might incur a fitness cost for maintaining long telomeres, because replication forks slow down at telomeres and can be stalled by Rap1 and secondary structures such as G-quadruplexes (Douglas and Diffley, 2021; Ivessa et al., 2002; Lopes et al., 2011; Paeschke et al., 2011), although no fitness defect was detected in a laboratory strain carrying longer telomeres (Harari et al., 2017). Keeping these two plausible selection forces in mind, our finding that the shortest 1-3% of telomeres in all strains are maintained at a roughly similar length, with only a relatively low correlation with the median length, would be consistent with selection acting only on the shortest telomeres to keep them just above a detrimental threshold. The 1^st^ percentile measured at ∼138 bp would provide some margin for the shortest telomeres to shorten and be elongated without crossing the proposed critical threshold lengths between 20 and 80 bp (Abdallah et al., 2009; Kockler et al., 2021; Rat et al., 2025; Sholes et al., 2022; Strecker et al., 2017). Accordingly, we show that natural strains with much longer telomeres generally display steady-state distributions that are more spread, hence the observed large CVs, rather than tight distributions with increased means. Remarkably, the shortest telomeres in a strain do not belong preferentially to one or a few defined extremities; rather, telomeres from any extremity have a similar probability of being among the shortest ones, thus spreading the potential risk associated with being short.

To conclude, we combine the most resolutive TL measurement method to date with a large sample set representing the phylogenetic and ecological diversity of *S. cerevisiae* to characterize the variability in TL distributions, its sources, and infer the effects of ploidy and subtelomere composition on TL. Our work therefore establishes a solid foundation for future studies of TL regulation harnessing the natural diversity of *S. cerevisiae*.

## Material and Methods

### Strains and sequencing data

We used data from 100 sequenced strains presented in (O’Donnell et al., 2023) (Supp. Table S1). In addition, strain AHG was sequenced again to test the reproducibility of our method, as were the WT W303 (*Mat***a** *ura3-1 trp1-1 leu2-3,112 his3-11,15 can1-100 RAD5 ADE2*) and *yku80Δ* (strain name: yZX440) strains. *YKU80* was deleted by replacement with *LEU2* using plasmid pFA6a-*LEU2* (Houseley and Tollervey, 2011) and standard PCR-based replacement methods (Longtine et al., 1998). The strains were grown in yeast extract, peptone, dextrose media (YPD) at 30°C.

### Artificial polyploid construction

Haploid and homozygous diploid derivatives of the North American (NA) YPS128 and West African (WA) DBVPG6044 strains (Cubillos et al., 2009; Louvel et al., 2014) were engineered to generate a default *a*-type haploid (YPB001 for the NA strain and YPB002 for the WA strain) and an *alpha*-mating–proficient diploid (YPB003 for the NA strain and YPB004 for the WA strain) (Supp. Table S1). Gene replacements at the *MAT* loci by *MATa-URA3* or *MATalpha-LEU2* were performed using standard lithium acetate transformation (Gietz and Woods, 2002). Strains were recursively mated to increase ploidy and stabilized using a functional isogenic *MATa* strain. Ploidy of parental and derived lines was verified by propidium iodide staining and flow cytometry (Supp. Fig. S3A), as described in (Gómez-Muñoz and Fischer, 2025).

### Genomic DNA extraction

We grew *S. cerevisiae* cells picked from a single colony in 10 mL YPD media at 30 °C overnight. A total number of cells of 7 × 10^9^ or fewer was used for DNA extraction. High-molecular-weight (HMW) DNA was extracted using QIAGEN Genomic-tip 100/G according to the manufacturer’s instructions. Alternatively, genomic DNA was extracted using the CTAB method, as described in (Kaur et al., 2018). Briefly, 2 × 10^9^ cells were incubated for 20 min at 37°C in spheroblasting solution (1 M sorbitol, 50 mM KPO_4_, 10 mM EDTA) with 1.25 mg of zymolyase 20T. Spheroblasts were pelleted and resuspended in CTAB extraction solution (3% CTAB, 100 mM Tris-HCl, 25 mM EDTA, 2 M NaCl and 2% PVP40) with 12.5 µL of 20 mg/mL proteinase K and 2 µL of 100 mg/mL RNase A. After 40 min of incubation at 37°C, the cellular debris were separated from DNA using chloroform:isoamyl alcohol solution. The CTAB-DNA mixture was subsequently precipitated using a CTAB dilution solution (1% CTAB, 50 mM Tris-HCl, 10 mM EDTA). The precipitate was washed with NaCl solution 1 (0.4 M NaCl, 10 mM Tris-HCl, 1 mM EDTA) to remove the excess of CTAB and then resuspended in NaCl solution 2 (1.42 M NaCl, 10 mM Tris-HCl, 1 mM EDTA). The DNA was precipitated by adding 2 volumes of absolute ethanol and centrifuging for 2 min at 12,000 rpm. After a wash with 70% ethanol, the DNA were resuspended in TE buffer. DNA quantity was controlled by Qubit dsDNA HS Assay.

### Oxford Nanopore Technologies sequencing

Library preparation and ONT sequencing were performed based on the protocol of ‘1D Native barcoding genomic DNA with EXP-NBD104 and SQK-LSK108’ when using FLO-MIN106 (R9.4.1) MinION flow cells and protocol of ‘1D Genomic DNA by Ligation with EXP-NBD104 and SQK-LSK109—PromethION’ when using the FLO-PRO114M (R10.4.1) flow cell. DNA repair and end preparation were performed on up to 2 μg of HMW DNA per sample using the NEBNext FFPE DNA Repair Mix with the following reaction setup: 48 μL DNA, 3.5 μL NEBNext FFPE DNA Repair Buffer, 2 μL NEBNext FFPE DNA Repair Mix, 3.5 μL Ultra II End Prep Reaction Buffer and 3 μL Ultra II End Prep Enzyme Mix, incubated at 20 °C for 15 min followed by 65 °C for 15 min. DNA size selection was then carried out using AMPure XP Beads (1:1 ratio) followed by native barcode ligation (22.5 μL DNA, 2.5 μL native barcode provided by EXP-NBD104 kit and 25 μL Blunt/TA Ligase Master Mix; at 25 °C for 20 min). After another round of AMPure XP bead clean-up (1:1 ratio), the samples were pooled together and adaptors were ligated at 25 °C for 15 min (65 μL DNA, 5 μL adapter mix II (AMII) provided by the EXP-NBD 104 kit, 20 μL NEBNext Quick Ligation Reaction Buffer and 10 μL Quick T4 DNA Ligase; at 25 °C for 15 min). The adaptor-ligated DNA sample was cleaned up by adding a 0.4× volume of AMPure XP beads followed by incubation for 5 min at room temperature. When using the SQK-LSK108 kit for FLO-MIN106 MinION flow cells, two successive 140-μL adapter bead binding buffer (ABB) washes were performed. When using the SQK-LSK109 kit for FLO-PRO114M flow cells, two successive 250-μL L fragment buffer washes were performed. The final library was eluted in 15 μL Elution Buffer and loaded into the MinION or PromethION flow cells according to the ONT manuals.

### Telomere length measurement pipeline

All bioinformatic analyses and pipelines were coded in Python (version 3.9.12) with Numpy (version 1.21.6) and Pandas (version 1.4.2) libraries installed. Raw fast5 files generated from sequencing were basecalled using the Super Accurate model of Guppy (version 6.4.2) followed by Porechop (version 0.2.4; https://github.com/rrwick/Porechop) removal of adapters and barcodes (Fig. 1A). Reads were then mapped to the genome assembly of the same strain using minimap2 (https://github.com/lh3/minimap2) (Li, 2018) to retrieve the ones mapping to telomeric regions. A mapping quality filter of MAPQ ≥ 30 was applied, eliminating 15% and keeping 85% of the mapped telomeric reads across all samples. We then ran Telofinder (https://github.com/GillesFischerSorbonne/telofinder) (O’Donnell et al., 2023) on these selected reads to detect the terminal telomeric sequences. Telofinder calculates the sequence entropy and proportions of dinucleotides (TG/CA, GG/CC and GT/AC) within a 20-bp sliding window, enabling the detection of degenerated *S. cerevisiae* telomere motifs (TG)_1-6_TG_2-3_/C_2-3_A(CA)_1-6_ on ONT reads when specific thresholds are met (polynuc_threshold = 0.8 and entropy_threshold = 0.8). To ensure that the identified sequences corresponded to true telomeres, we verified that a telomeric sequence found at the beginning (5’-end) of a read was C-rich, and that one found at the end (3’-end) was G-rich, as expected based on the 5’-3’ orientation of sequencing through the pores. Additionally, for G-rich telomere sequences found at 3’ ends, we selected only reads that contained a terminal adapter to ensure that they were not truncated before sequencing reached the end of the DNA molecule.

### Phylogenetic tree processing and visualization

The phylogenetic tree (Fig. 2A) was built based on data from (O’Donnell et al., 2023), with adjustments to retain only the 100 leaves representing the 100 strains analyzed here. We used the Bio.Phylo package from Biopython (v1.79) for phylogenetic analysis and iTOL v6 (Letunic and Bork, 2024) for visualization.

### Plots and statistical analyses

Python with Seaborn (version 0.11.2) and SciPy (version 1.7.3) libraries was used to draw plots and perform the statistical analyses.

### Data access

All raw and processed sequencing data (fast5, fastq and fasta files) generated in this study have been deposited to the European Nucleotide Archive (https://www.ebi.ac.uk/ena/ browser/home) under the project PRJEB89249.

### Code availability

All the scripts used in this publication can be found as Supplementary Code and are also available at https://github.com/Telomere-Genome-Stability/Yeast2025.

## Acknowledgement

We thank members of GF’s and ZX’s laboratories for discussions. Research in ZX’s laboratory was supported by Agence Nationale de la Recherche (ANR-24-CE12-7740-01 and ANR-24-CE12-0083-03) and Institut National du Cancer (PLBIO20-312). Research in GF’s laboratory was supported by Agence Nationale de la Recherche (ANR-24-CE12-7740-02 and ANR-24-CE12-0998-03). EK’s work at the Biomics Platform, C2RT, Institut Pasteur, Paris, France, was supported by France Génomique (ANR-10-INBS-09).

## Author contributions

CG: Methodology, Investigation, Software, Formal Analysis, Writing – Original Draft. CGM: Methodology, Investigation. EK: Methodology, Investigation, Software. NA: Investigation. OI: Investigation. GF: Conceptualization, Methodology, Supervision, Writing – Original Draft. ZX: Investigation, Conceptualization, Supervision, Writing – Original Draft. All authors: Writing – Review & Editing.

## References

Abdallah, P., Luciano, P., Runge, K.W., Lisby, M., Geli, V., Gilson, E., and Teixeira, M.T. (2009). A two-step model for senescence triggered by a single critically short telomere. Nat Cell Biol 11, 988–993.

Allsopp, R.C., Vaziri, H., Patterson, C., Goldstein, S., Younglai, E.V., Futcher, A.B., Greider, C.W., and Harley, C.B. (1992). Telomere length predicts replicative capacity of human fibroblasts. Proc. Natl. Acad. Sci. U. S. A. 89, 10114–10118.

Blackburn, E.H., and Chiou, S.S. (1981). Non-nucleosomal packaging of a tandemly repeated DNA sequence at termini of extrachromosomal DNA coding for rRNA in Tetrahymena. Proc. Natl. Acad. Sci. U. S. A. 78, 2263–2267.

Cristofari, G., and Lingner, J. (2006). Telomere length homeostasis requires that telomerase levels are limiting. EMBO J.

Cubillos, F.A., Louis, E.J., and Liti, G. (2009). Generation of a large set of genetically tractable haploid and diploid Saccharomyces strains. FEMS Yeast Res 9, 1217–1225.

D’Angiolo, M., Yue, J.X., De Chiara, M., Barre, B.P., Giraud Panis, M.J., Gilson, E., and Liti, G. (2023). Telomeres are shorter in wild Saccharomyces cerevisiae isolates than in domesticated ones. Genetics 223.

de Lange, T. (2018). Shelterin-Mediated Telomere Protection. Annu. Rev. Genet. 52, 223–247.

Douglas, M.E., and Diffley, J.F.X. (2021). Budding yeast Rap1, but not telomeric DNA, is inhibitory for multiple stages of DNA replication in vitro. Nucleic Acids Res 49, 5671–5683.

Eberhard, S., Valuchova, S., Ravat, J., Fulnecek, J., Jolivet, P., Bujaldon, S., Lemaire, S.D., Wollman, F.A., Teixeira, M.T., Riha, K., et al. (2019). Molecular characterization of Chlamydomonas reinhardtii telomeres and telomerase mutants. Life science alliance 2.

Enomoto, S., Glowczewski, L., and Berman, J. (2002). MEC3, MEC1, and DDC2 Are Essential Components of a Telomere Checkpoint Pathway Required for Cell Cycle Arrest during Senescence in Saccharomyces cerevisiae. Mol. Biol. Cell. 13, 2626–2638.

Fajkus, J., Kovarik, A., Kralovics, R., and Bezdek, M. (1995). Organization of telomeric and subtelomeric chromatin in the higher plant Nicotiana tabacum. Mol. Gen. Genet. 247, 633–638.

Fulcher, N., Teubenbacher, A., Kerdaffrec, E., Farlow, A., Nordborg, M., and Riha, K. (2015). Genetic architecture of natural variation of telomere length in Arabidopsis thaliana. Genetics 199, 625–635.

Gallardo, F., Laterreur, N., Cusanelli, E., Ouenzar, F., Querido, E., Wellinger, R.J., and Chartrand, P. (2011). Live cell imaging of telomerase RNA dynamics reveals cell cycle-dependent clustering of telomerase at elongating telomeres. Mol. Cell 44, 819–827.

Gatbonton, T., Imbesi, M., Nelson, M., Akey, J.M., Ruderfer, D.M., Kruglyak, L., Simon, J.A., and Bedalov, A. (2006). Telomere Length as a Quantitative Trait: Genome-Wide Survey and Genetic Mapping of Telomere Length-Control Genes in Yeast. PLoS Genet 2, e35.

Gietz, R.D., and Woods, R.A. (2002). Transformation of yeast by lithium acetate/single-stranded carrier DNA/polyethylene glycol method. Methods Enzymol. 350, 87–96.

Gomes, N.M., Ryder, O.A., Houck, M.L., Charter, S.J., Walker, W., Forsyth, N.R., Austad, S.N., Venditti, C., Pagel, M., Shay, J.W., et al. (2011). Comparative Biology of Mammalian Telomeres: Hypotheses on Ancestral States and the Roles of Telomeres in Longevity Determination. Aging Cell.

Gómez-Muñoz, C., and Fischer, G. (2025). MuPETFlow: multiple ploidy estimation tool from flow cytometry data. BMC Genomics 26, 301.

Greider, C.W., and Blackburn, E.H. (1985). Identification of a specific telomere terminal transferase activity in Tetrahymena extracts. Cell 43, 405–413.

Harari, Y., Zadok-Laviel, S., and Kupiec, M. (2017). Long Telomeres Do Not Affect Cellular Fitness in Yeast. mBio 8.

Houseley, J., and Tollervey, D. (2011). Repeat expansion in the budding yeast ribosomal DNA can occur independently of the canonical homologous recombination machinery. Nucleic Acids Res 39, 8778–8791.

Hu, C., Zhu, X.T., He, M.H., Shao, Y., Qin, Z., Wu, Z.J., and Zhou, J.Q. (2024). Elimination of subtelomeric repeat sequences exerts little effect on telomere essential functions in Saccharomyces cerevisiae. eLife 12.

Ijpma, A.S., and Greider, C.W. (2003). Short Telomeres Induce a DNA Damage Response in Saccharomyces cerevisiae. Mol. Biol. Cell 14, 987–1001.

Ivessa, A.S., Zhou, J.Q., Schulz, V.P., Monson, E.K., and Zakian, V.A. (2002). Saccharomyces Rrm3p, a 5’ to 3’ DNA helicase that promotes replication fork progression through telomeric and subtelomeric DNA. Genes Dev. 16, 1383–1396.

Jain, D., and Cooper, J.P. (2010). Telomeric strategies: means to an end. Annual Review of Genetics 44, 243–269.

Karimian, K., Groot, A., Huso, V., Kahidi, R., Tan, K.T., Sholes, S., Keener, R., McDyer, J.F., Alder, J.K., Li, H., et al. (2024). Human telomere length is chromosome end-specific and conserved across individuals. Science 384, 533–539.

Kaur, H., Ahuja, J.S., and Lichten, M. (2018). Methods for Controlled Protein Depletion to Study Protein Function during Meiosis. Methods Enzymol. 601, 331–357.

Kockler, Z.W., Comeron, J.M., and Malkova, A. (2021). A unified alternative telomere-lengthening pathway in yeast survivor cells. Mol. Cell.

Letunic, I., and Bork, P. (2024). Interactive Tree of Life (iTOL) v6: recent updates to the phylogenetic tree display and annotation tool. Nucleic Acids Res 52, W78–W82.

Levy, D.L., and Blackburn, E.H. (2004). Counting of Rif1p and Rif2p on Saccharomyces cerevisiae telomeres regulates telomere length. Mol. Cell. Biol. 24, 10857–10867.

Li, H. (2018). Minimap2: pairwise alignment for nucleotide sequences. Bioinformatics 34, 3094–3100.

Liti, G., Haricharan, S., Cubillos, F.A., Tierney, A.L., Sharp, S., Bertuch, A.A., Parts, L., Bailes, E., and Louis, E.J. (2009). Segregating YKU80 and TLC1 alleles underlying natural variation in telomere properties in wild yeast. PLoS Genet 5, e1000659.

Longtine, M.S., McKenzie, A.I., Demarini, D.J., Shah, N.G., Wach, A., Brachat, A., Philippsen, P., and Pringle, J.R. (1998). Additional modules for versatile and economical PCR-based gene deletion and modification in Saccharomyces cerevisiae. Yeast 14, 953–961.

Lopes, J., Piazza, A., Bermejo, R., Kriegsman, B., Colosio, A., Teulade-Fichou, M.P., Foiani, M., and Nicolas, A. (2011). G-quadruplex-induced instability during leading-strand replication. EMBO J. 30, 4033–4046.

Louvel, H., Gillet-Markowska, A., Liti, G., and Fischer, G. (2014). A set of genetically diverged Saccharomyces cerevisiae strains with markerless deletions of multiple auxotrophic genes. Yeast 31, 91–101.

Lundblad, V., and Szostak, J.W. (1989). A mutant with a defect in telomere elongation leads to senescence in yeast. Cell 57, 633–643.

Marcand, S., Gilson, E., and Shore, D. (1997). A protein-counting mechanism for telomere length regulation in yeast. Science 275, 986–990.

Monaghan, P., Eisenberg, D.T.A., Harrington, L., and Nussey, D. (2018). Understanding diversity in telomere dynamics. Philos. Trans. R. Soc. Lond. B. Biol. Sci. 373.

Moyzis, R.K., Buckingham, J.M., Cram, L.S., Dani, M., Deaven, L.L., Jones, M.D., Meyne, J., Ratliff, R.L., and Wu, J.R. (1988). A highly conserved repetitive DNA sequence, (TTAGGG)n, present at the telomeres of human chromosomes. Proc. Natl. Acad. Sci. U. S. A. 85, 6622–6626.

Mozdy, A.D., and Cech, T.R. (2006). Low abundance of telomerase in yeast: implications for telomerase haploinsufficiency. RNA 12, 1721–1737.

O’Donnell, S., Yue, J.X., Saada, O.A., Agier, N., Caradec, C., Cokelaer, T., De Chiara, M., Delmas, S., Dutreux, F., Fournier, T., et al. (2023). Telomere-to-telomere assemblies of 142 strains characterize the genome structural landscape in Saccharomyces cerevisiae. Nat. Genet. 55, 1390–1399.

Paeschke, K., Capra, J.A., and Zakian, V.A. (2011). DNA Replication through G-Quadruplex Motifs Is Promoted by the Saccharomyces cerevisiae Pif1 DNA Helicase. Cell 145, 678–691.

Peter, J., De Chiara, M., Friedrich, A., Yue, J.X., Pflieger, D., Bergstrom, A., Sigwalt, A., Barre, B., Freel, K., Llored, A., et al. (2018). Genome evolution across 1,011 Saccharomyces cerevisiae isolates. Nature 556, 339–344.

Porter, S.E., Greenwell, P.W., Ritchie, K.B., and Petes, T.D. (1996). The DNA-binding protein Hdf1p (a putative Ku homologue) is required for maintaining normal telomere length in Saccharomyces cerevisiae. Nucleic Acids Res 24, 582–585.

Rat, A., Martinez Fernandez, V., Doumic, M., Teixeira, M.T., and Xu, Z. (2025). Mathematical model linking telomeres to senescence in Saccharomyces cerevisiae reveals cell lineage versus population dynamics. Nat Commun 16, 1024.

Sanchez, S.E., Gu, Y., Wang, Y., Golla, A., Martin, A., Shomali, W., Hockemeyer, D., Savage, S.A., and Artandi, S.E. (2024). Digital telomere measurement by long-read sequencing distinguishes healthy aging from disease. Nat Commun 15, 5148.

Schmidt, T.T., Tyer, C., Rughani, P., Haggblom, C., Jones, J.R., Dai, X., Frazer, K.A., Gage, F.H., Juul, S., Hickey, S., et al. (2024). High resolution long-read telomere sequencing reveals dynamic mechanisms in aging and cancer. Nat Commun 15, 5149.

Shakirov, E.V., and Shippen, D.E. (2004). Length regulation and dynamics of individual telomere tracts in wild-type Arabidopsis. Plant Cell 16, 1959–1967.

Shampay, J., and Blackburn, E.H. (1988). Generation of telomere-length heterogeneity in Saccharomyces cerevisiae. Proc. Natl. Acad. Sci. U. S. A. 85, 534–538.

Sholes, S.L., Karimian, K., Gershman, A., Kelly, T.J., Timp, W., and Greider, C.W. (2022). Chromosome-specific telomere lengths and the minimal functional telomere revealed by nanopore sequencing. Genome Res. 32, 616-628.

Strecker, J., Stinus, S., Caballero, M.P., Szilard, R.K., Chang, M., and Durocher, D. (2017). A sharp Pif1-dependent threshold separates DNA double-strand breaks from critically short telomeres. eLife 6.

Teixeira, M.T. (2013). Saccharomyces cerevisiae as a Model to Study Replicative Senescence Triggered by Telomere Shortening. Front Oncol 3, 101.

Teixeira, M.T., Arneric, M., Sperisen, P., and Lingner, J. (2004). Telomere Length Homeostasis Is Achieved via a Switch between Telomerase-Extendible and -Nonextendible States. Cell 117, 323–335.

Teplitz, G.M., Pasquier, E., Bonnell, E., De Laurentiis, E., Bartle, L., Lucier, J.F., Sholes, S., Greider, C.W., and Wellinger, R.J. (2024). A mechanism for telomere-specific telomere length regulation. bioRxiv.

Walmsley, R.M., and Petes, T.D. (1985). Genetic control of chromosome length in yeast. Proc. Natl. Acad. Sci. U. S. A. 82, 506–510.

Wellinger, R.J., and Zakian, V.A. (2012). Everything you ever wanted to know about Saccharomyces cerevisiae telomeres: beginning to end. Genetics 191, 1073–1105.

Wicky, C., Villeneuve, A.M., Lauper, N., Codourey, L., Tobler, H., and Muller, F. (1996). Telomeric repeats (TTAGGC)n are sufficient for chromosome capping function in Caenorhabditis elegans. Proc. Natl. Acad. Sci. U. S. A. 93, 8983–8988.

Xu, Z., Dao Duc, K., Holcman, D., and Teixeira, M.T. (2013). The length of the shortest telomere as the major determinant of the onset of replicative senescence. Genetics 194, 847–857.

