## Supplementary Information for "Natural diversity of telomere length distributions across 100 *Saccharomyces cerevisiae* strains"

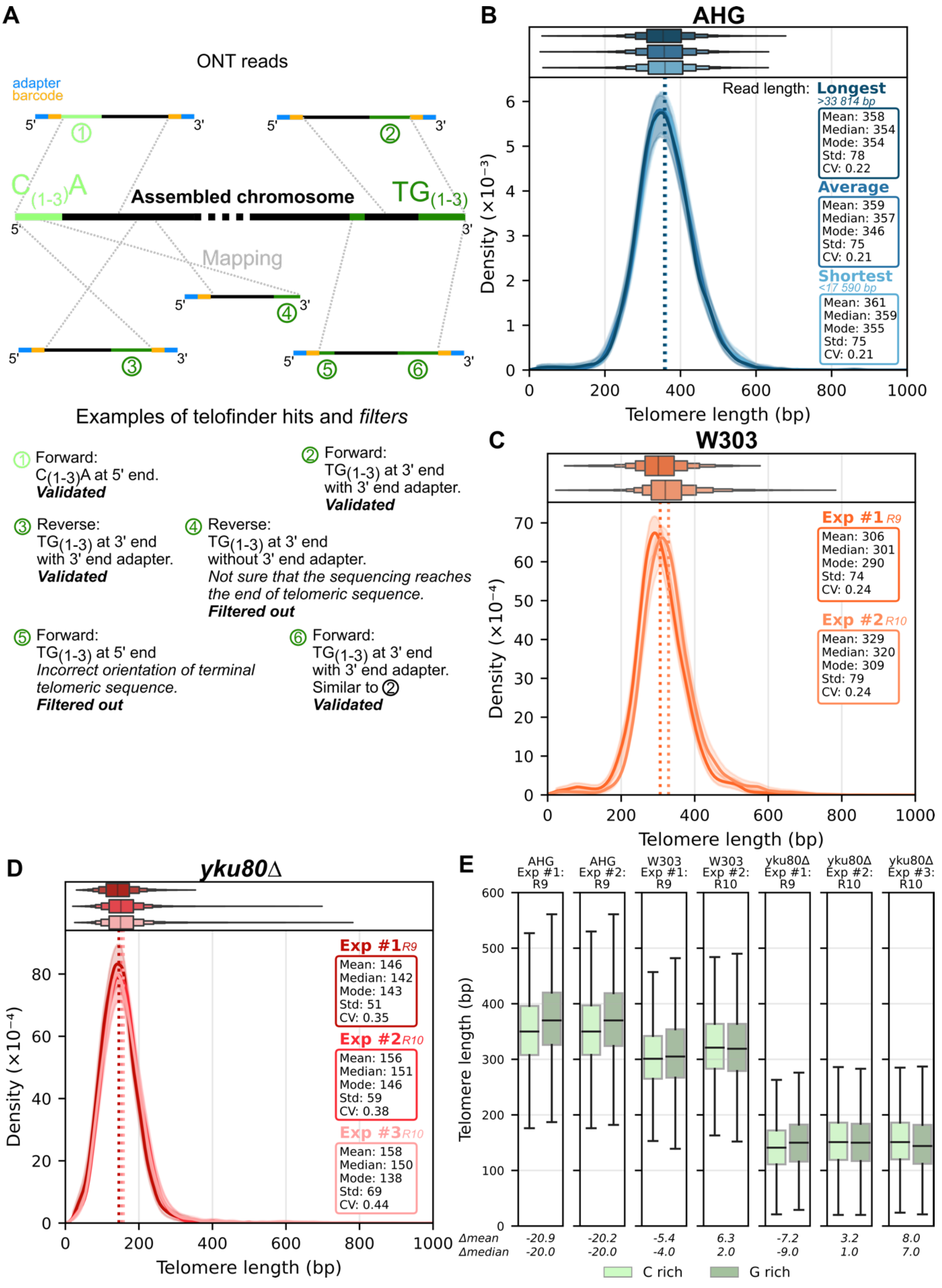


**Supplementary Figure 1. Robust TL distribution measurement from Nanopore reads.** (A) Filters used to eliminate telomere-like sequences with the wrong orientation and internal telomere sequences, and to ensure the read reached the end the molecule. Examples numbered 1 to 6 illustrate validated or filtered out reads. (B) TL distributions computed from equally sized subsets of reads of different lengths (“Longest” >34 kb, “Average” between 18 and 34 kb, and “Shortest” <18 kb), using the AHG reads from (O'Donnell et al., 2023). (C) Independent biological replicates of W303’s TL distribution. (D) Independent biological replicates of the *yku80Δ* mutant’s TL distribution. (E) Comparison between TL distributions derived from the C-rich or G-rich strands for the indicated experiments. (C-E) The version of the ONT flowcell technology used, “R9” (for R9.4.1) or “R10” (for R10.4.1), is indicated.

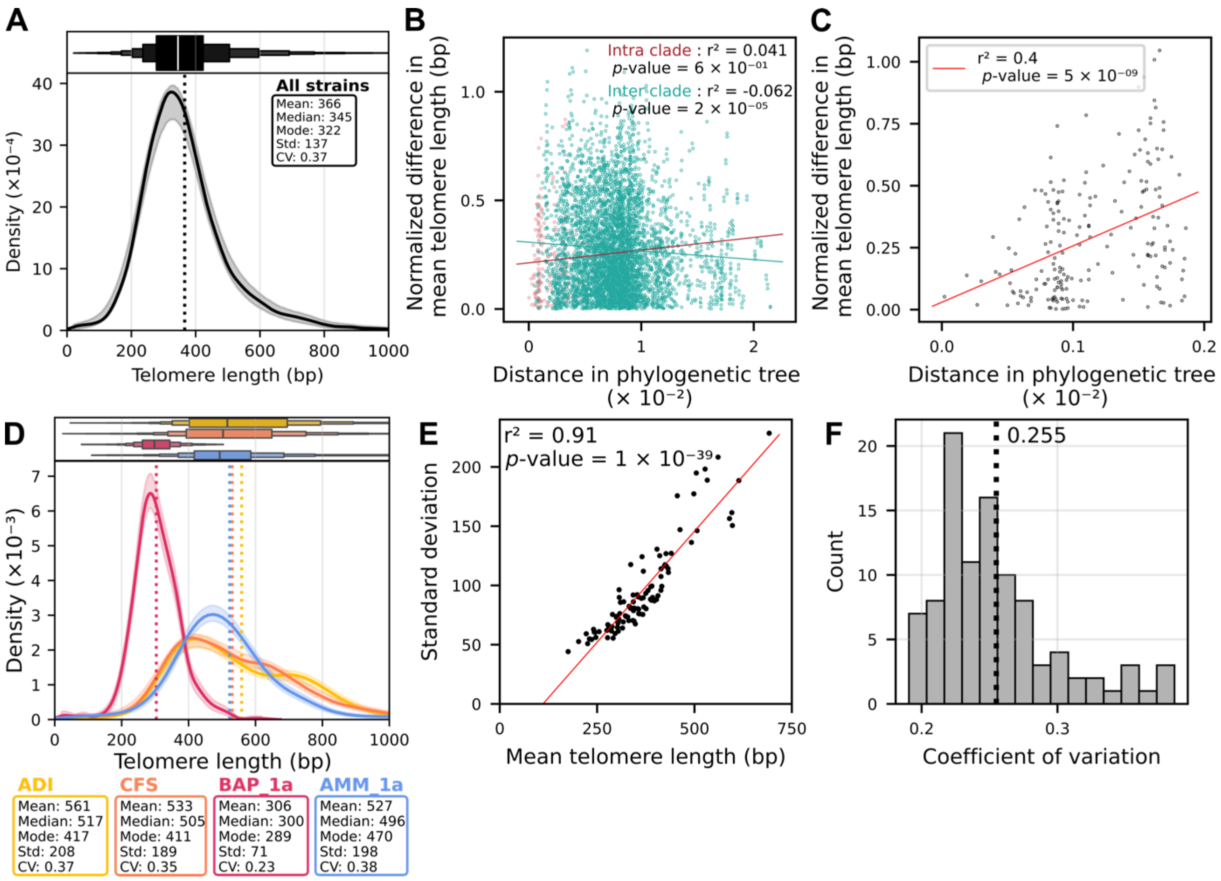


**Supplementary Figure 2. Characterization of TL distribution metrics in natural strains.** (A) Representative overview of TL distribution in the species computed from a balanced random sample of telomeric reads from each strain. (B) Correlation between phylogenetic distance and the normalized difference in mean TL (difference between the 2 TL means divided by the average of the 2 means). Each dot represents a pair of strains. A pair within the same clade or between 2 clades is shown in red or cyan, respectively. Pearson's correlation ($r^2$) and associated *p*-values are shown. (C) Same as (B), with a focus on the 4% closest pairs of strains. (D) TL distributions for strains ADI, CFS, BAP_1a and AMM_1a. (E) Correlation between mean TL and standard deviation across strains. (F) Distribution of the CVs of all strains.

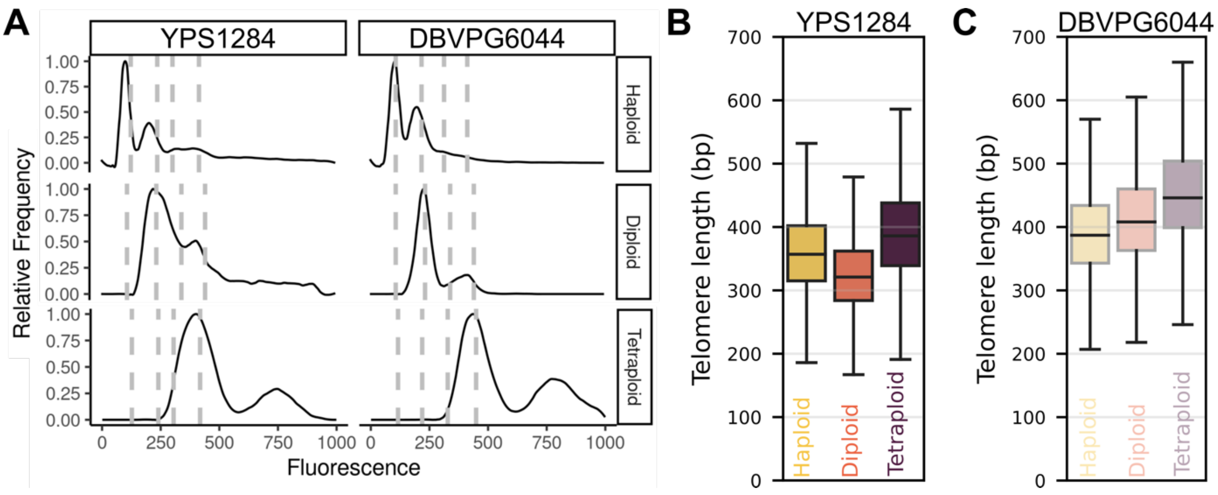


**Supplementary Figure 3. Tetraploids exhibit longer telomeres than isogenic diploids and haploids.** (A) Propidium iodide staining and flow cytometry analysis of DNA content of the indicated haploid, diploid, and tetraploid strains. Vertical dotted lines represent 1C, 2C, 3C, and 4C DNA contents based on positive control strains used in the same flow cytometry experiment. (B) Boxplots of global TL distributions of isogenic haploid, diploid, and tetraploid strains from the same West African DBVPG6044 strain background. (C) Same as (B) with the North American YPS128 strain background.

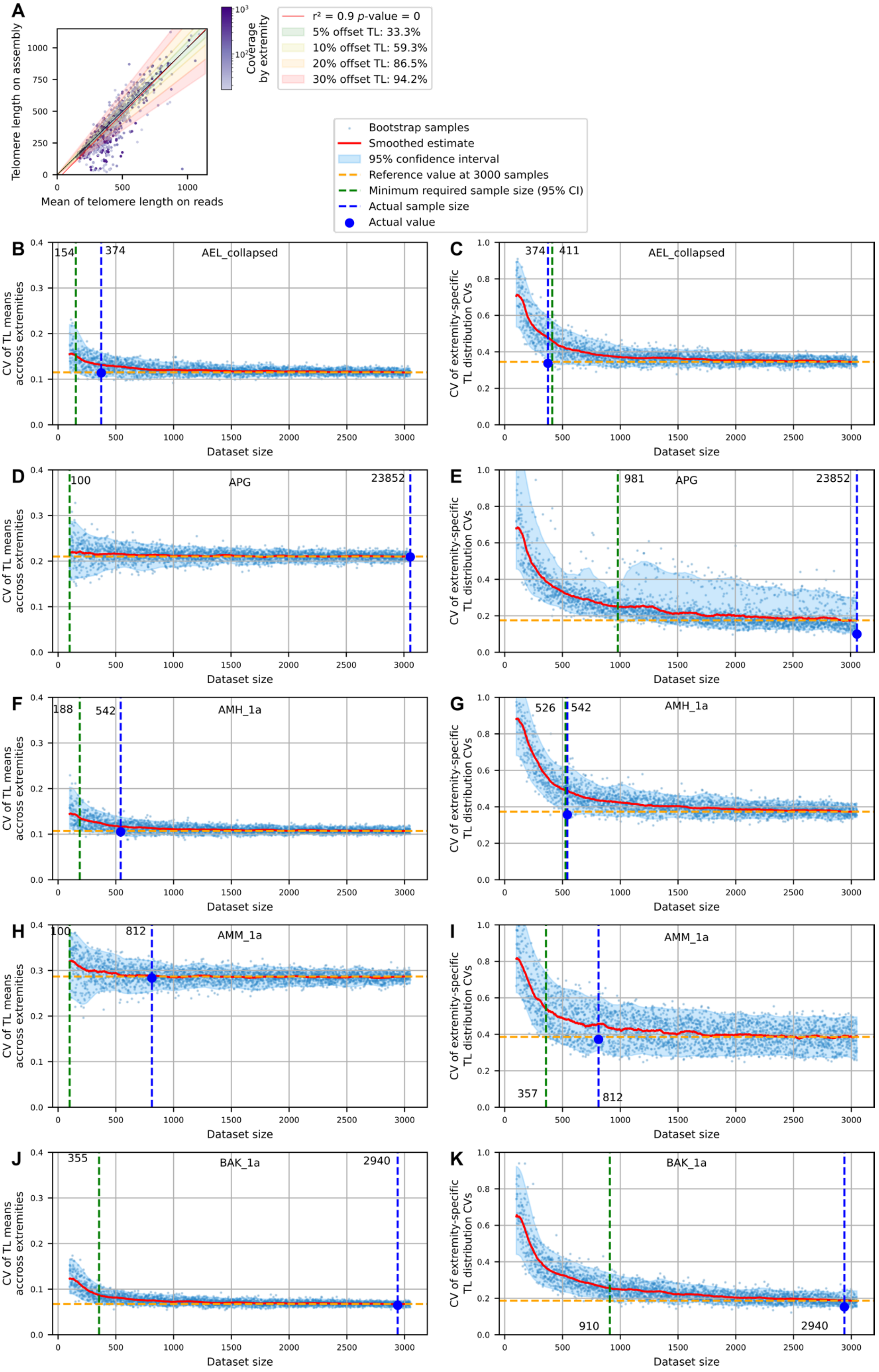
A
Telomere length on assembly
Mean of telomere length on reads
Coverage by extremity
r² = 0.9 p-value = 0
5% offset TL: 33.3%
10% offset TL: 59.3%
20% offset TL: 86.5%
30% offset TL: 94.2%
Bootstrap samples
Smoothed estimate
95% confidence interval
Reference value at 3000 samples
Minimum required sample size (95% CI)
Actual sample size
Actual value
B
AEL_collapsed
CV of TL means accross extremities
Dataset size
C
CV of extremity-specific TL distribution CVs
D
APG
E
F
AMH_1a
G
H
AMM_1a
I
J
BAK_1a
K

**Supplementary Figure 4. Comparison between assembly-derived and read-derived extremity-specific TL values, and sample size requirements.** (A) Extremity-specific TL values derived from genome assemblies from (O'Donnell et al., 2023) and means of extremity-specific TL distributions computed from reads are plotted. Each dot represents the telomere at one extremity and its read coverage is indicated by its color. Shaded areas of different colors display 5-30% offsets between the two values and the percentage in the legend indicate the fraction of affected extremities. (B, D, F, H, J) Bootstrap analyses of the CV of extremity-specific TL means as a function of the size of the dataset (number of telomeric reads). Each blue dot corresponds to one bootstrap simulation. The average is represented by the red line and the shaded blue area corresponds to 95% of the bootstrap values. The horizontal yellow dotted line shows the value for a dataset size of 3000. The green dotted line indicates the threshold sample size, corresponding to the point where the blue area diverges from the yellow line. The blue dotted line shows the actual dataset size. We determine that the sample size is adequate if the green dotted line is on the left of the blue dotted line. (C, E, G, I, K) Same as (B, D, F, H, J) but for the CV of the extremity-specific TL distribution CVs. (B) and (C) show the example of a strain that does not meet the criteria for the CV of the extremity-specific TL distribution CVs. (D-K) shows this analysis for the 4 strains displayed in Fig. 4A.

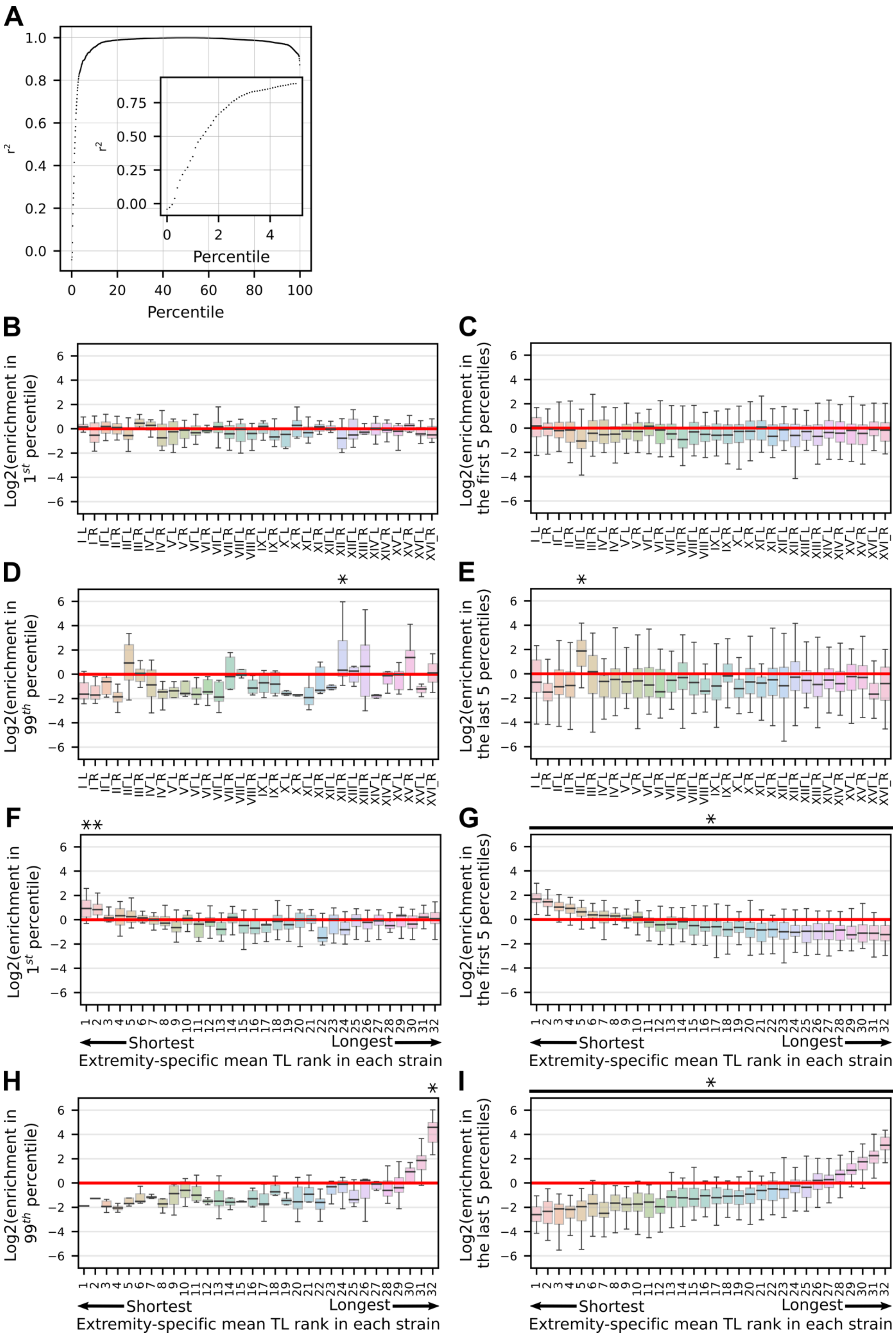
A
r²
Percentile
B
Log2(enrichment in 1st percentile)
C
Log2(enrichment in the first 5 percentiles)
D
Log2(enrichment in 99th percentile)
E
Log2(enrichment in the last 5 percentiles)
F
Log2(enrichment in 1st percentile)
G
Log2(enrichment in the first 5 percentiles)
H
Log2(enrichment in 99th percentile)
I
Log2(enrichment in the last 5 percentiles)
I_L I_R II_L II_R III_L III_R IV_L IV_R V_L V_R VI_L VI_R VII_L VII_R VIII_L VIII_R IX_L IX_R X_L X_R XI_L XI_R XII_L XII_R XIII_L XIII_R XIV_L XIV_R XV_L XV_R XVI_L XVI_R
Shortest
Longest
Extremity-specific mean TL rank in each strain

**Supplementary Figure 5. Contributions of specific extremities to the shortest or longest telomeres.** (A) Coefficient of determination $r^2$ between the median TL and the $n^{th}$ percentile (x-axis). Inset shows a zoom on the first 5 percentile. (B) Enrichment of specific chromosome extremities in the set of individual telomeres shorter than the $1^{st}$ percentile in a strain-specific TL distribution. The boxplot represents the enrichment values from all strains in Log2 scale. (C) Same as (B) for the $5^{th}$ percentile. (D) and (E), same as (B) and (C) but for the set of individual telomeres longer than the $99^{th}$ (resp. $95^{th}$) percentile. (F) Enrichment of the chromosome extremities ranked for each strain according to their mean extremity-specific TL in the set of individual telomeres shorter than the $1^{st}$ percentile. (G) Same as (F) for the $5^{th}$ percentile. (H) and (I), same as (G) and (F) but for the set of individual telomeres longer than the $99^{th}$ (resp. $95^{th}$) percentile. (B-I) In these panels, an ANOVA test was performed to detect differences in means. The following *p*-values were obtained, respectively: 0.28, 0.069, 0.034, $1.8x10^{-15}$, 0.0065, $9.8x10^{-129}$, $2.2x10^{-17}$, $9.6x10^{-177}$. * indicates extremities that are significantly different from at least two others based on *p*-values from Tukey's HSD tests with a 0.05 threshold.

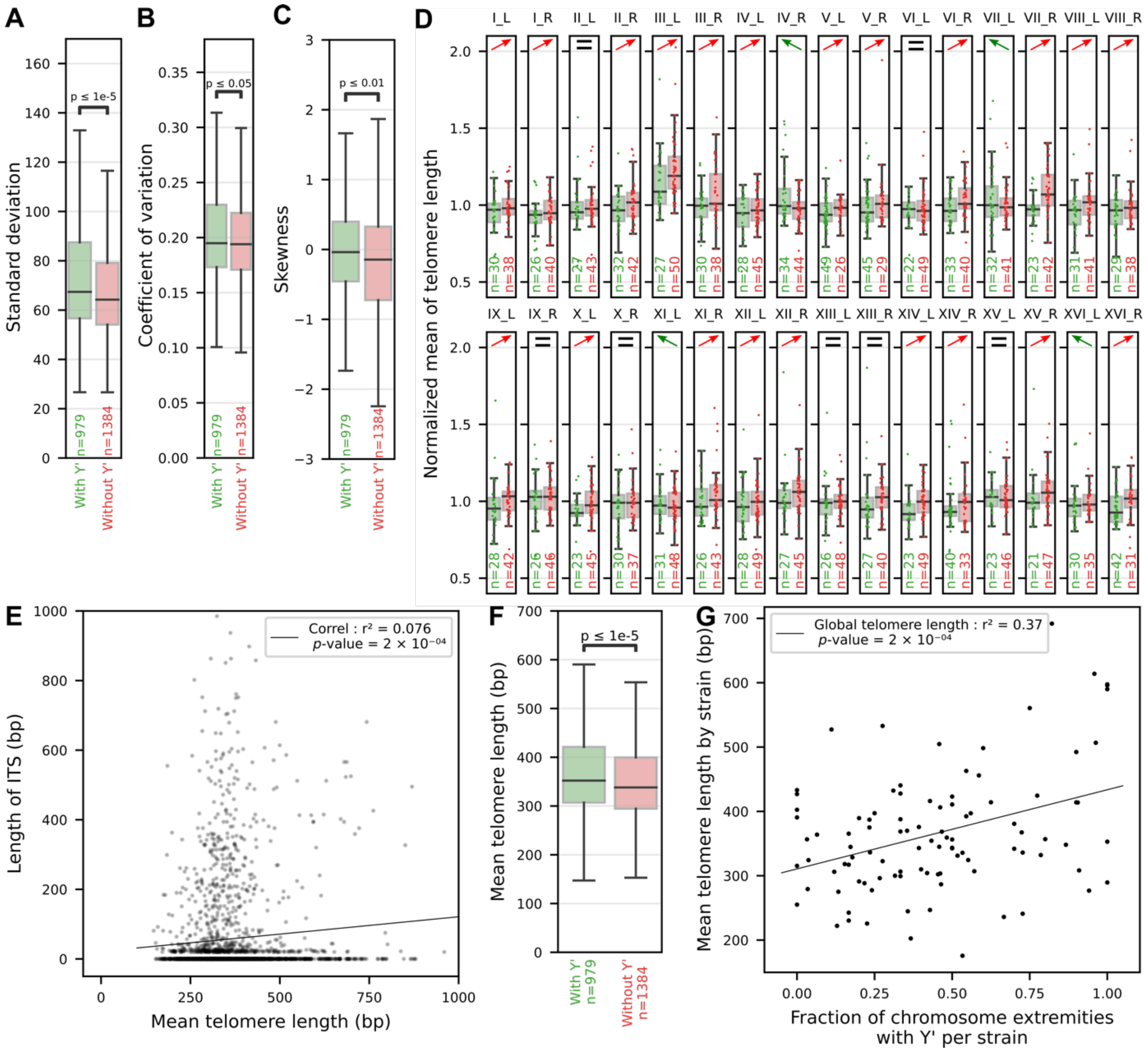


**Supplementary Figure 6. Correlation between the presence of a Y' element and TL.** (A) Standard deviations of the extremity-specific TL distributions, clustered depending on the presence or absence of Y' element in the corresponding subtelomere. The number of chromosome extremities in each group is indicated by "n". The *p*-value from a Student's t-test is indicated. (B) Same as (A) but for the coefficient of variation of the extremity-specific TL distribution. (C) Same as (A) but for the skewness. (D) Distributions of the normalized extremity-specific TL means shown for each extremity across strains, clustered depending on the presence or absence of Y' element in the corresponding subtelomere. Red and green arrow indicate the direction of the differences in TL means, while "=" indicates identical means within a margin of 1%. (E) Plot representing the total length of ITSs at a chromosome extremity against the mean TL at the same extremity, across all strains, showing no correlation. (F) Distributions of extremity-specific TL means across the extremities of all strains, clustered according to the presence or absence of the Y' element in the corresponding subtelomere. (G) Correlation between the global TL means and the fraction of extremities with Y' elements in the corresponding strain.

**Supplementary Table S1. List of strains and accessions.**

Summary table of all strains used in this work. Relevant sequencing statistics and TL distribution metrics are indicated.

**Supplementary Code. Codes used in this work to measure telomere length from ONT sequencing data.**

### References

O'Donnell, S., Yue, J.X., Saada, O.A., Agier, N., Caradec, C., Cokelaer, T., De Chiara, M., Delmas, S., Dutreux, F., Fournier, T.*, et al.* (2023). Telomere-to-telomere assemblies of 142 strains characterize the genome structural landscape in Saccharomyces cerevisiae. Nat. Genet. *55*, 1390-1399.
