## Supplementary Code for "Natural diversity of telomere length distributions across 100 *Saccharomyces cerevisiae* strains": README.md

For an up-to-date version, please visit https://github.com/Telomere-Genome-Stability/Yeast2025.

# Yeast2025

Code for pipeline presented in the paper 'Natural diversity of telomere length distributions across 100 *Saccharomyces cerevisiae* strains' Clotilde Garrido, Cintia Gómez-Muñoz,Etienne Kornobis, Nicolas Agier, Oana Ilioaia, Gilles Fischer, Zhou Xu.

@author: Clotilde Garrido, Sorbonne Université - CNRS

A Python pipeline for detecting and extracting telomeric regions from long-read (ONT) sequencing data in *Saccharomyces cerevisiae*.

# Features

- Telomere detection on assembled genomes
- Read mapping with Minimap2
- Region-based read extraction
- Adapter trimming analysis
- Telofinder re-analysis on extracted reads
- Read quality filtering (MAPQ, terminal location, adapter presence)
- Generates final per-strain, per-region summary tables

# Dependencies

Install required tools:

- Telofinder (https://telofinder.readthedocs.io/en/latest/#installation)
- minimap 2 (https://github.com/lh3/minimap2)
- samtools (https://github.com/samtools/samtools)
- seqkit (https://bioinf.shenwei.me/seqkit/#installation)
  
# Configuration

Before running the pipeline, you need to specify the full path to the `telofinder.py` script.

Edit the configuration variable `TELOFINDER_PATH` in the `run_pipeline.py` script to point to the correct location:

```python
TELOFINDER_PATH = '/full/path/to/telofinder.py'
```

# Usage

```bash
python run_pipeline.py \
    --input_data data/example_input.csv \
    --output_dir results/ \
    --threads 8 \
    --intern_min_length 100 \
    --only_trim
```

# Arguments

| Argument                   | Description                                                                    | Default value  |
|----------------------------|--------------------------------------------------------------------------------|----------------|
| `-i`, `--input_data`       | Input TSV file containing file paths                                           | required       |
| `-o`, `--output_dir`       | Output directory for the results.                                              | required       |
| `-t`, `--threads`          | Number of threads to use.                                                      | 3              |
| `-l`, `--intern_min_length`|Keep internal telomeric assemblies if their length is above this value.         | `no`           |
| `-trim`, `--only_trim`     | Use only already trimmed reads; the original untrimmed reads are not available.| `false`        |

## Input Table Format

The pipeline requires an input TSV file listing all strains and their associated files. This table must include the following columns:

| Column name    | Description                                                                                         |
|----------------|-----------------------------------------------------------------------------------------------------|
| `Strain`       | Unique identifier for each yeast strain.                                                            |
| `Reference`    | Path to the assembled genome file in FASTA format. This file is required.                           |
| `Trim_reads`   | Path to adapter-trimmed long-read sequencing data in FASTA format.                                  |
| `Notrim_reads` | Path to raw (untrimmed) long-read sequencing data in FASTA format. Optional if using `--only_trim`. |

**Example input TSV (tab-separated):**

```tsv
Strain	Reference	Trim_reads	Notrim_reads
Strain1	/path/to/genome.fasta	/path/to/trimmed.fastq	/path/to/raw.fastq
Strain2	/path/to/genome2.fasta	/path/to/trimmed2.fastq	/path/to/raw2.fastq
```

# Outputs

All output files and folders are generated inside the directory specified by the `--output_dir` argument. The pipeline produces the following outputs:

- `Telofinder_Assembly/`  
  Telomere detection results on assembled genomes.

- `Mapping/`  
  Sorted BAM files resulting from mapping trimmed reads to the reference genome.

- `Extract_reads/`  
  Extracted reads corresponding to specific telomeric regions.

- `Telofinder_Reads/`  
  Telofinder re-analysis results on the extracted reads.

- `telofinder_result_merged_assembly.csv`  
  Summary table combining telomere calls from all regions and strains.

- `All_Results.csv`  
  Table containing all telomeric stretches found in reads.

- `Filtred_Results.csv`  
  Final filtered table containing high-quality telomeric reads after all quality controls.
