## Supplementary Code for "Natural diversity of telomere length distributions across 100 *Saccharomyces cerevisiae* strains": Run_pipeline.py

```
# -*- coding: utf-8 -*-

# === USER CONFIGURATION REQUIRED ===
# Please provide the full path to the telofinder.py file you downloaded or cloned.
TELOFINDER_PATH = '/path/to/telofinder.py'


# === DO NOT MODIFY BELOW THIS LINE UNLESS NEEDED ===
import os
import subprocess
import pandas as pd 
import numpy as np
import argparse
import sys

def parse_arguments():
    parser = argparse.ArgumentParser(description="Pipeline for detecting terminal telomeres in nanopore reads.")
    parser.add_argument('-i','--input_data', type=str, required=True, help="Input TSV file containing file paths")
    parser.add_argument('-o','--output_dir', type=str, required=True, help="Output directory for the results.")
    parser.add_argument('-t', '--threads', type=str, default='3', help="Number of threads to use. Default is 3.")
    parser.add_argument('-l', '--intern_min_length', type=str, default='no', help="Keep internal telomeric assemblies if their length is above this value. Default is 'no'")
    parser.add_argument('-trim', '--only_trim', action='store_true', help="Use only already trimmed reads; the original untrimmed reads are not available.")
    return parser.parse_args()

def check_csv_file(csv_path,trim):
    """
    Validates the input TSV file.
    
    Parameters:
        csv_path (str): Path to the input TSV file.
        trim (bool): Whether only trimmed reads are provided (no access to untrimmed reads).
    
    Returns:
        pandas.DataFrame: The validated TSV content as a DataFrame.
    """

    # Load the TSV file
    try:
        data = pd.read_csv(csv_path,sep='\t')
    except Exception as e:
        sys.exit(f"Unable to load the TSV file: {e}")

    # Define required columns
    required_columns = ["Strain", "Reference", "Notrim_reads", "Trim_reads"]
    if trim == True:
        required_columns = ["Strain", "Reference", "Trim_reads"]
    # Check required columns
    if not all(col in data.columns for col in required_columns):
        print(f"Available columns: {list(data.columns)}")
        sys.exit(f"The CSV file must contain the following columns: {', '.join(required_columns)}")

    # 1. Ensure 'Strain' values are unique
    if data["Strain"].duplicated().any():
        duplicates = data[data["Strain"].duplicated(keep=False)]
        sys.exit(f"The 'Strain' column contains duplicate entries:\n{duplicates['Strain'].tolist()}.\n"
            f"Please rename them (e.g., AAB_1, AAB_2, etc.).")

    # 2. Check for missing or invalid reference files
    if data["Reference"].isna().any():
        missing_reference_rows = data[data["Reference"].isna()]
        sys.exit(f"Some rows are missing a reference file. Please check the following:\n{missing_reference_rows}")
    for ref in data["Reference"].dropna().unique():
        if not os.path.exists(ref):
            sys.exit(f"The reference file '{ref}' does not exist. Please verify the path.")

    # 3. Validate read file paths
    reads_col=["Notrim_reads", "Trim_reads"]
    if trim == True:
        reads_col=[ "Trim_reads"]
    for col in reads_col:
    	# Ensure no missing values
        if data[col].isna().any():
            missing_rows = data[data[col].isna()]
            sys.exit(f"Some rows in column '{col}' are empty. Please check the following:\n{missing_rows}")
        # Ensure files exist
        for file_path in data[col].dropna():
            if not os.path.exists(file_path):
                sys.exit(f"The file '{file_path}' in column '{col}' does not exist.")
    print("* The CSV file is valid.\n")
    return(data)

def run_telofinder_on_assembly(assembly_path, ref, output_dir,threads):
    """
    Run Telofinder on a genome assembly.
    
    Parameters:
        assembly_path (str): Path to the genome assembly file.
        ref (str): Identifier for the assembly (used in output naming).
        output_dir (str): Directory to store Telofinder results.
        threads (str): Number of threads to use.
    """
    cmd = f'python {TELOFINDER_PATH} --force -o {output_dir}Telofinder_Assembly/telofinder_result_{ref}_assembly -t {threads} {assembly_path}'
    print(f"* Running Telofinder on assembly {ref} : {cmd}\n")
    subprocess.run(cmd, shell=True, check=True,stdout=subprocess.DEVNULL,stderr=subprocess.DEVNULL)

def merged_telofinder_on_assembly(data, intern_min_len, output_dir): 
    """ 
       Merge and filter Telofinder results from multiple assemblies.

    Parameters:
        data (DataFrame): Input data table containing references.
        intern_min_len (str): Minimum length for internal telomeres to keep (or 'no' to skip).
        output_dir (str): Directory where Telofinder outputs are located.

    Returns:
        DataFrame: Merged and filtered Telofinder results.
    """
    TFa = pd.DataFrame()
    for assembly_path in list(set(data['Reference'])):
        ref = assembly_path.split('/')[-1].rsplit(".", 1)[0]
        tf=pd.read_csv(f'{output_dir}Telofinder_Assembly/telofinder_result_{ref}_assembly/merged_telom_df.csv', sep=',').dropna()
        for i in tf.index:
            if tf.at[i,'type']=='term' or (intern_min_len !='no' and tf.at[i,'len'] > int(intern_min_len)):
                ind=len(TFa)
                TFa.at[ind,'assembly']=ref
                TFa.at[ind,'chrom']=tf.at[i,'chrom']
                TFa.at[ind,'side']=tf.at[i,'side']
                TFa.at[ind,'type']=tf.at[i,'type']
                TFa.at[ind,'start']=tf.at[i,'start']
                TFa.at[ind,'end']=tf.at[i,'end']
                TFa.at[ind,'len']=tf.at[i,'len']
                TFa.at[ind,'chrom_size']=tf.at[i,'chrom_size']
    TFa.to_csv(f'{output_dir}telofinder_result_merged_assembly.csv', sep='\t', index=False)
    return(TFa)

def map_reads_to_assembly(assembly_path, porechop_path, label, output_dir,threads):
    """
    Map trimmed reads to the genome assembly using Minimap2, then sort and index the resulting BAM.

    Parameters:
        assembly_path (str): Path to the genome assembly file (FASTA).
        trimmed_reads_path (str): Path to the trimmed reads (FASTA).
        label (str): Label used to name output files.
        output_dir (str): Directory to store the BAM output.
        threads (str): Number of threads to use.
    """
    bam_output = f'{output_dir}{label}.sorted.bam'
    cmd = f'minimap2 -a -x map-ont {assembly_path} {porechop_path} -K 5M -t {threads} | samtools sort --threads {threads} > {bam_output}'
    print(f"* Mapping reads with Minimap2 {label} : {cmd}\n")
    subprocess.run(cmd, shell=True, check=True,stdout=subprocess.DEVNULL,stderr=subprocess.DEVNULL)
    subprocess.run(f'samtools index -@ {threads} {bam_output}', shell=True, check=True,stdout=subprocess.DEVNULL,stderr=subprocess.DEVNULL)

def extract_reads_from_bam(bam_file, notrim_file, chrom, start, end, output_dir, strain):
    """
    Extract reads mapped to a specific region from a BAM file and convert them to FASTA format.
    If untrimmed reads are available, also extract the corresponding sequences from them.

    Parameters:
        bam_file (str): Path to the BAM file.
        notrim_file (str): Path to the original untrimmed reads (or 'No' if unavailable).
        chrom (str): Chromosome or contig name.
        start (int): Start position of the region.
        end (int): End position of the region.
        output_dir (str): Output directory for extracted reads.
        strain (str): Sample identifier.

    Returns:
        int: Number of extracted reads.
    """
    ext_bam = f'{output_dir}Extract_reads/{strain}-{chrom}_{start}-{end}.bam'
    ext_sam = f'{output_dir}Extract_reads/{strain}-{chrom}_{start}-{end}.sam'
    ext_fasta = f'{output_dir}Extract_reads/{strain}-{chrom}_{start}-{end}.fasta'
    if notrim_file != 'No':
        ext_notrim_fasta = f'{output_dir}Extract_reads/{strain}-{chrom}_{start}-{end}_notrim.fasta'
    cmd_bam = f'samtools view -h {bam_file} {chrom}:{start}-{end} -b -o {ext_bam}'
    cmd_fasta = f'samtools fasta {ext_bam} > {ext_fasta}'
    cmd_sam = f'samtools view -h {ext_bam} > {ext_sam}'
    if notrim_file != 'No':
        cmd_seq = f'seqkit seq -n -i {ext_fasta} > {output_dir}/tmp.txt'
        cmd_grep = f'seqkit grep -f {output_dir}tmp.txt {notrim_file} -o {ext_notrim_fasta}'
        cmd_count = f'grep ">" -c {ext_notrim_fasta} > {output_dir}tmp.txt'
    else:
        cmd_count = f'grep ">" -c {ext_fasta} > {output_dir}tmp.txt'
    subprocess.run(cmd_bam, shell=True, check=True,stdout=subprocess.DEVNULL,stderr=subprocess.DEVNULL)
    subprocess.run(cmd_fasta, shell=True, check=True,stdout=subprocess.DEVNULL,stderr=subprocess.DEVNULL)
    subprocess.run(cmd_sam, shell=True, check=True,stdout=subprocess.DEVNULL,stderr=subprocess.DEVNULL)
    if notrim_file != 'No':
        subprocess.run(cmd_seq, shell=True, check=True,stdout=subprocess.DEVNULL,stderr=subprocess.DEVNULL)
        subprocess.run(cmd_grep, shell=True, check=True,stdout=subprocess.DEVNULL,stderr=subprocess.DEVNULL)
    try:
        subprocess.run(cmd_count, shell=True, check=True,stdout=subprocess.DEVNULL,stderr=subprocess.DEVNULL)
        with open(f"{output_dir}tmp.txt", "r") as f:
            nb = int(f.read().strip())
    except subprocess.CalledProcessError as e:
        if e.returncode == 1:
            nb = 0
    return(nb)

def run_telofinder_on_reads(fasta_file, out_dir, threads):
    """
    Run Telofinder on extracted reads in FASTA format.

    Parameters:
        fasta_file (str): Path to the FASTA file containing reads.
        out_dir (str): Output directory for Telofinder results.
        threads (str or int): Number of threads to use.
    """
    cmd = f'python {TELOFINDER_PATH} --force -o {out_dir} -t {threads} {fasta_file}'
    subprocess.run(cmd, shell=True, check=True,stdout=subprocess.DEVNULL,stderr=subprocess.DEVNULL)

def load_fasta(file_path):
    """
    Load sequences from a FASTA file into a dictionary.

    Parameters:
        file_path (str): Path to the FASTA file.

    Returns:
        dict: Dictionary where keys are sequence headers and values are sequences.
    """
    sequences = {}
    with open(file_path, 'r') as file:
        header = ''
        sequence = ''
        for line in file:
            line = line.strip()
            if line.startswith('>'):
                if header != '':
                    sequences[header] = sequence
                header = line[1:].split()[0]
                sequence = ''
            else:
                sequence += line
        sequences[header] = sequence
    return sequences


def check_adapter_presence(original_file, modified_file):
    """
    Compare original and trimmed FASTA reads to determine if adapter sequences were removed.

    Parameters:
        original_file (str): Path to the original (untrimmed) FASTA file.
        modified_file (str): Path to the modified (trimmed) FASTA file.

    Returns:
        DataFrame: Summary of adapter presence at the start and end of each read.
    """
    DF=pd.DataFrame()
    original_sequences = load_fasta(original_file)
    modified_sequences = load_fasta(modified_file)
    for header, original_seq in original_sequences.items():
        i=len(DF)
        DF.at[i,'read']=header
        modified_seq = modified_sequences.get(header, '')
        if len(modified_seq) == 0:
            DF.at[i,'check']='Split'
        else:
            if len(original_seq.split(modified_seq)[0])==0:
                DF.at[i,'check']='NO start adapt'
                DF.at[i,'seq start adapt']=''
                DF.at[i,'len start adapt']=0
                DF.at[i,'size_trim_read']=len(modified_seq)
            else :
                DF.at[i,'check']='start adapt'
                DF.at[i,'seq start adapt']=original_seq.split(modified_seq)[0]
                DF.at[i,'len start adapt']=len(DF.at[i,'seq start adapt'])
                DF.at[i,'size_trim_read']=len(modified_seq)
            if len(original_seq.split(modified_seq)[1])==0:
                DF.at[i,'check']+='_NO end adapt'
                DF.at[i,'seq end adapt']=''
                DF.at[i,'len end adapt']=0
                DF.at[i,'size_trim_read']=len(modified_seq)
            else:
                DF.at[i,'check']+='_end adapt'
                DF.at[i,'seq end adapt']=original_seq.split(modified_seq)[1]
                DF.at[i,'len end adapt']=len(DF.at[i,'seq end adapt'])
                DF.at[i,'size_trim_read']=len(modified_seq)
                # Debug: unexpected case where trimmed sequence is not found in original
                if modified_seq not in original_seq : 
                    print(f"Warning: trimmed sequence not found in original for read '{header}'")
    return(DF)


def main(args):
    """
    Main function to execute the full pipeline with given arguments.
    """

    if not os.path.isfile(TELOFINDER_PATH):
        raise FileNotFoundError(f"The file {TELOFINDER_PATH} does not exist. Please update TELOFINDER_PATH with the correct location.")
    trim = args.only_trim
    data = check_csv_file(args.input_data,trim)
    output_dir = args.output_dir
    T = args.threads
    intern_min_len = args.intern_min_length

    # Step 1: Run Telofinder on reference assemblies
    if 'Telofinder_Assembly' not in os.listdir(output_dir):
         os.mkdir(output_dir+'Telofinder_Assembly')
    for assembly_path in list(set(data['Reference'])):
         ref = assembly_path.split('/')[-1].rsplit(".", 1)[0]
         run_telofinder_on_assembly(assembly_path, ref, output_dir,T)
    TFa = merged_telofinder_on_assembly(data, intern_min_len, output_dir)
    print(f"Step 1 completed: Telofinder results on assemblies are saved in {output_dir}Telofinder_Assembly/")

    # Step 2: Map trimmed reads to assemblies using Minimap2
    if 'Mapping' not in os.listdir(output_dir):
        os.mkdir(output_dir+'Mapping')
    for _, row in data.iterrows():
        assembly_path = row['Reference']
        ref = assembly_path.split('/')[-1].rsplit(".", 1)[0]
        label = ref+'_'+row['Strain']
        trim_path = row['Trim_reads']
        map_reads_to_assembly(assembly_path, trim_path, label, output_dir+'Mapping/',T)
    print(f"Step 2 completed: Reads mapping BAM files are saved in {output_dir}Mapping/")

    # Step 3: Extract reads from BAM files and convert to FASTA
    if 'Extract_reads' not in os.listdir(output_dir):
        os.mkdir(output_dir+'Extract_reads')
    for _, row in data.iterrows():
        strain = row['Strain']
        ref = row['Reference'].split('/')[-1].rsplit(".", 1)[0]
        label = ref+'_'+strain
        bam_file = f'{output_dir}Mapping/{label}.sorted.bam'
        if trim == False:
            reads_file = row['Notrim_reads']
        else:
            reads_file = 'No'
        print(f"* Extracting reads for {strain}\n")
        for i in TFa[TFa['assembly']==ref].index:
            chrom = TFa.at[i,'chrom']
            start=str(int(TFa.at[i,'start']))
            end=str(int(TFa.at[i,'end']))
            nb = extract_reads_from_bam(bam_file, reads_file, chrom, start, end, output_dir, strain)
            TFa.at[i,'#reads_'+strain]=nb
    TFa.to_csv(f'{output_dir}telofinder_result_merged_assembly.csv', sep='\t', index=False)
    print(f"Step 3 completed: Extracted reads FASTA and BAM files are saved in {output_dir}Extract_reads/")

    # Step 4: Run Telofinder again on the extracted reads
    if 'Telofinder_Reads' not in os.listdir(output_dir):
         os.mkdir(output_dir+'Telofinder_Reads')
    for _, row in data.iterrows():
        strain = row['Strain']
        ref = row['Reference'].split('/')[-1].rsplit(".", 1)[0]
        label = ref+'_'+strain
        print(f"* Running Telofinder on reads extracted for {strain}\n")
        for i in TFa[TFa['assembly']==ref].index:
            if TFa.at[i,'#reads_'+strain]>0:
                chrom = TFa.at[i,'chrom']
                start=str(int(TFa.at[i,'start']))
                end=str(int(TFa.at[i,'end']))
                if trim == False:
                    fasta_file = f'{output_dir}Extract_reads/{strain}-{chrom}_{start}-{end}_notrim.fasta'
                else:
                    fasta_file = f'{output_dir}Extract_reads/{strain}-{chrom}_{start}-{end}.fasta'
                out_dir = f'{output_dir}Telofinder_Reads/{strain}-{chrom}_{start}-{end}'
                run_telofinder_on_reads(fasta_file, out_dir,T)
    print(f"Step 4 completed: Telofinder results on extracted reads are saved in {output_dir}Telofinder_Reads/")

    # Step 5: Collect Telofinder results
    Res=pd.DataFrame()
    for _, row in data.iterrows():
        strain = row['Strain']
        print(f"* Collecting results for  {strain}\n")
        ref = row['Reference'].split('/')[-1].rsplit(".", 1)[0]
        for i in TFa[TFa['assembly']==ref].index:
            if TFa.at[i,'#reads_'+strain]>0:
                chrom = TFa.at[i,'chrom']
                start=str(int(TFa.at[i,'start']))
                end=str(int(TFa.at[i,'end']))
                if trim == False : 
                    fasta_file = f'{output_dir}Extract_reads/{strain}-{chrom}_{start}-{end}_notrim.fasta'
                    trim_file = f'{output_dir}Extract_reads/{strain}-{chrom}_{start}-{end}.fasta'
                    adapt = check_adapter_presence(fasta_file, trim_file)
                sam_file = f'{output_dir}Extract_reads/{strain}-{chrom}_{start}-{end}.sam'
                Sam = pd.read_csv(sam_file,delim_whitespace=True,names=['QNAME','FLAG','RNAME','POS','MAPQ','CIGAR','RNEXT','PNEXT','TLEN','SEQ','QUAL','NM','ms','AS','nn','tp','cm','s1','s2','dv','SA'], index_col=False,comment='@', engine="python")
                result_file = f'{output_dir}Telofinder_Reads/{strain}-{chrom}_{start}-{end}/merged_telom_df.csv'
                df = pd.read_csv(result_file, sep=',').dropna()

                for read in set(df['chrom']):
                    samr=Sam[Sam['QNAME']==read]
                    if len(samr)>1:
                        samr=samr[samr['MAPQ']==max(samr['MAPQ'])]
                        if len(samr)>1:
                            samr=samr[samr['FLAG']==min(samr['FLAG'])]
                    ind_sam=list(samr.index)[0]
                    if trim == False :
                        ind_adapt=adapt[adapt['read']==read].index[0]
                    for j in df[df['chrom']==read].index:
                        ind = len(Res)
                        Res.at[ind,'assembly']=ref
                        Res.at[ind,'chrom']=chrom
                        Res.at[ind,'chrom_side']=TFa.at[i,'side']
                        Res.at[ind,'chrom_type']=TFa.at[i,'type']
                        Res.at[ind,'chrom_start']=start
                        Res.at[ind,'chrom_end']=end
                        Res.at[ind,'chrom_len']=TFa.at[i,'len']
                        Res.at[ind,'chrom_size']=TFa.at[i,'chrom_size']
                        Res.at[ind,'strain']=strain
                        Res.at[ind,'#reads']=TFa.at[i,'#reads_'+strain]
                        Res.at[ind,'read']=df.at[j,'chrom']
                        Res.at[ind,'side']=df.at[j,'side']
                        if trim == False :
                            if df.at[j,'type']=='term' or (df.at[j,'side']=='Left' and df.at[j,'start']-adapt.at[ind_adapt,'len start adapt']<20) or (df.at[j,'side']=='Right' and df.at[j,'chrom_size']-df.at[j,'end']-adapt.at[ind_adapt,'len end adapt']<20):
                            	Res.at[ind,'type']='term'
                            else:
                                Res.at[ind,'type']='intern'
                        else :
                            Res.at[ind,'type']=df.at[j,'type']
                        Res.at[ind,'start']=df.at[j,'start']
                        Res.at[ind,'end']=df.at[j,'end']
                        Res.at[ind,'len']=df.at[j,'len']
                        if trim == False :
                            Res.at[ind,'read_size_notrim']=df.at[j,'chrom_size']
                            Res.at[ind,'read_size_trim']=adapt.at[ind_adapt,'size_trim_read']
                        else :
                            Res.at[ind,'read_size_trim']=df.at[j,'chrom_size']

                        Res.at[ind,'MAPQ']=samr.at[ind_sam,'MAPQ']
                        Res.at[ind,'FLAG']=samr.at[ind_sam,'FLAG']
                        if trim == False :
                            Res.at[ind,'check']=adapt.at[ind_adapt,'check']
                            Res.at[ind,'seq start adapt']=adapt.at[ind_adapt,'seq start adapt']
                            Res.at[ind,'len start adapt']=adapt.at[ind_adapt,'len start adapt']
                            Res.at[ind,'seq end adapt']=adapt.at[ind_adapt,'seq end adapt']
                            Res.at[ind,'len end adapt']=adapt.at[ind_adapt,'len end adapt']

    Res.to_csv(f'{output_dir}All_Results.csv', sep='\t', index=False)
    print(f"Step 5 completed: All_Results.csv has been created at {output_dir}")

    # Step 6: Apply filters
    Res = Res[Res['chrom_type']=='term']
    Res = Res[Res['MAPQ']>=30] # MAPQ >= 30
    Res = Res[Res['type']=='term'] 
    ind=[]
    if trim == False :
        for i in Res.index:
            if Res.at[i,'side']=='Left': # Left = Crich
                if Res.at[i,'start']-Res.at[i,'len start adapt'] < 20 :
                    ind.append(i)
            else:
                if Res.at[i,'len end adapt'] > 0: #Adaptateur telo en fin de read 
                    if Res.at[i,'read_size_notrim']-Res.at[i,'end']-Res.at[i,'len end adapt']<20:#Right = Grich
                        ind.append(i)
    else:
        for i in Res.index:
            if Res.at[i,'side']=='Left': # Left = Crich
                if Res.at[i,'start'] < 20 :
                    ind.append(i)
            else:
                if Res.at[i,'read_size_trim']-Res.at[i,'end']<20:#Right = Grich
                    ind.append(i)

    Res = Res.loc[ind]
    Res.to_csv(f'{output_dir}Filtred_Results.csv', sep='\t', index=False)

    Res = pd.read_csv(f'{output_dir}Filtred_Results.csv', sep='\t')
    print(f"Step 6 completed: Filtred_Results.csv has been created at {output_dir}")

    # Step 7: Add filtered read count and mean length per region
    for _, row in data.iterrows():
        strain = row['Strain']
        ref = row['Reference'].split('/')[-1].rsplit(".", 1)[0]
        label = ref+'_'+strain
        for i in TFa[TFa['assembly']==ref].index:
            chrom = TFa.at[i,'chrom']
            start=int(TFa.at[i,'start'])
            end=int(TFa.at[i,'end'])
            res = Res[(Res['assembly']==ref)&(Res['chrom']==chrom)&(Res['chrom_start']==start)&(Res['chrom_end']==end)&(Res['strain']==strain)]
            nb = len(res)
            mean = np.mean(res['len'])
            TFa.at[i,'#reads_'+strain+'filtr']=nb
            TFa.at[i,'Mean_reads_'+strain+'filtr']=mean
    TFa.to_csv(f'{output_dir}telofinder_result_merged_assembly.csv', sep='\t', index=False)
    print(f"Step 7 completed: telofinder_result_merged_assembly.csv has been updated at {output_dir}")

if __name__ == "__main__":
    args = parse_arguments()
    main(args)
```
